# A modular cell-free protein biosensor platform using split T7 RNA polymerase

**DOI:** 10.1101/2024.07.19.604303

**Authors:** Megan A. McSweeney, Alexandra T. Patterson, Kathryn Loeffler, Regina Cuellar Lelo de Larrea, Monica P. McNerney, Ravi S. Kane, Mark P. Styczynski

## Abstract

Conventional laboratory protein detection techniques are not suitable for point-of-care (POC) use because they require expensive equipment and laborious protocols, and existing POC assays suffer from long development timescales. Here, we describe a modular cell-free biosensing platform for generalizable protein detection that we call TLISA (**T**7 RNA polymerase-**L**inked **I**mmuno**S**ensing **A**ssay), designed for extreme flexibility and equipment-free use. TLISA uses a split T7 RNA polymerase fused to affinity domains against a protein. The target antigen drives polymerase reassembly, inducing reporter expression. We characterize the platform, then demonstrate its modularity by using 16 affinity domains against four different antigens with minimal protocol optimization. We show TLISA is suitable for POC use by sensing human biomarkers in serum and saliva with a colorimetric readout within one hour and by demonstrating functionality after lyophilization. Altogether, this technology could have potentially revolutionary impacts, enabling truly rapid, reconfigurable, equipment-free detection of virtually any protein.

## Introduction

Diagnosis at the earliest manifestations of disease is often critical in disease management and treatment at both the individual patient and public health levels. Unfortunately, most gold standard clinical diagnostic tests—including high performance liquid chromatography (HPLC), mass spectrometry, reverse transcriptase polymerase chain reaction (RT-PCR), and enzyme linked immunosorbent assays (ELISA)—require expensive equipment and trained technicians working in a well-funded laboratory setting (1,2). The costly nature of these techniques impedes sufficiently broad public access, especially in low-resource areas with little access to high-quality healthcare and in many low-income countries where annual per capita healthcare expenditures are less than US$100(3). Point-of-care (POC) diagnostic tools have a substantial impact on the management of both individual and public health by increasing test accessibility, which enables earlier detection of disease, more ideal treatment plans, and better prognoses (2). However, existing POC platforms like the lateral flow assay (LFA) are not simple to reconfigure to new targets due to the complex, iterative optimization of multiple interconnected design parameters that is required (4,5). Thus, engineering rapid, effective, and affordable POC tools is a dire need.

Cell-free expression (CFE) systems are a particularly promising platform poised to address current needs for POC diagnostic technology. These systems are composed simply of cellular lysate, substrates, and energy sources needed to execute transcription and translation from DNA templates *in vitro*. CFE systems can be lyophilized for long-term storage at ambient temperature (6), react robustly in small volumes of human sample matrices (7–10), can yield colorimetric outputs for equipment-free result interpretation (11), and cost only $0.02-$0.04 per 1 μL reaction (6). Numerous CFE biosensors have been engineered to detect nucleic acid (6,8,12–14), ion (9,11,15,16), and small molecule (7,10,16–18) biomarkers, with particularly impressive modularity and versatility in detecting nucleic acid sequences.

However, these technologies neglect an important class of biomarker: proteins, which are gold standard biomarkers for many diseases and the target of over 100 FDA-approved diagnostic tests (19). Despite this critical importance, our abilities for modular detection of proteins using CFE are far less mature.

Only a few efforts for modular protein sensing using CFE have been reported, all of which use aptamers (oligonucleotides that bind to a specific target with high affinity) to regulate gene expression upon protein recognition. Modular implementation of newly evolved or synthetic aptamers remains a challenge, though, with integration into existing sensing platforms typically done through costly and time-consuming trial-and-error experimentation (20). A recent plug-and-play DNA aptamer platform sought to address this issue in an *in vitro* transcription system, but relies on the aptamers forming a specific noncanonical DNA structure called a G-quadruplex, limiting the sensing space of the platform significantly (21). A new automated design process was also recently reported that converts arbitrary RNA aptamers into riboswitches for use in CFE biosensors (22). While this is a significant advance in the long-standing challenge of riboswitch engineering, it requires users to know the aptamer’s secondary structure when bound to the antigen and the antigen’s free energy of binding, specifications that are particularly challenging for new protein targets and thus impede its more widespread use. Critically, there currently are no aptamer-based CFE biosensors that have been linked to visual colorimetric outputs, tested in human samples, or lyophilized for storage and transportation, rendering these systems unsuitable for POC use.

To address this gap in capabilities and in the literature, we devised a strategy combining small, target-specific protein-binding affinity domains with a split T7 RNA polymerase (T7RNAP) to create a modular, easily engineerable, field-deployable protein detection platform. While antibodies are widely used in laboratory assays (e.g., ELISA) and in POC LFAs as specific and sensitive affinity domains, they are challenging to implement efficiently in CFE biosensing platforms due to post-translational modifications and challenges in proper folding (23,24). Thus, we chose to focus on the use of nanobodies (single-chain camelid immunoglobulin fragments), which can be expressed easily in bacterial CFE systems (25). Nanobodies are easy to engineer because they are small, highly stable, and incredibly robust to changes in their chemical environment and to fusions with diverse molecules (26). Moreover, nanobodies can be rapidly evolved to bind to new targets through processes such as phage, yeast, or ribosome display (27,28), in contrast to the more expensive and complex development pipeline for new antibody development (29). The split T7RNAP we use has one fragment that was evolved (T7RNAP_Nev_) to have minimal spontaneous reassembly in the absence of forced co-localization; that co-localization can be caused by binding to a target via affinity domains fused to the polymerase fragments (30,31). To our knowledge, neither of these approaches have been used for cell-free biosensing despite their broad utility having been demonstrated in *in vivo* applications.

By fusing nanobodies or other affinity domains to each of the split T7RNAP fragments, we have created a biosensing platform we call TLISA (**T**7RNAP-**L**inked **I**mmuno**S**ensing **A**ssay), analogous to an ELISA but in a faster, more user-friendly, and easily engineerable format (**Fig. 1A**). In the presence of a target antigen, both nanobodies bind, driving reassembly of the polymerase fragments and inducing expression of a reporter protein. Here, we demonstrate the modularity and utility of the TLISA platform, characterizing design considerations using eGFP and mCherry as model target antigens and then showing that the modular nature of TLISA enables rapid sensor development for clinically relevant proteins.

**Fig. 1:**
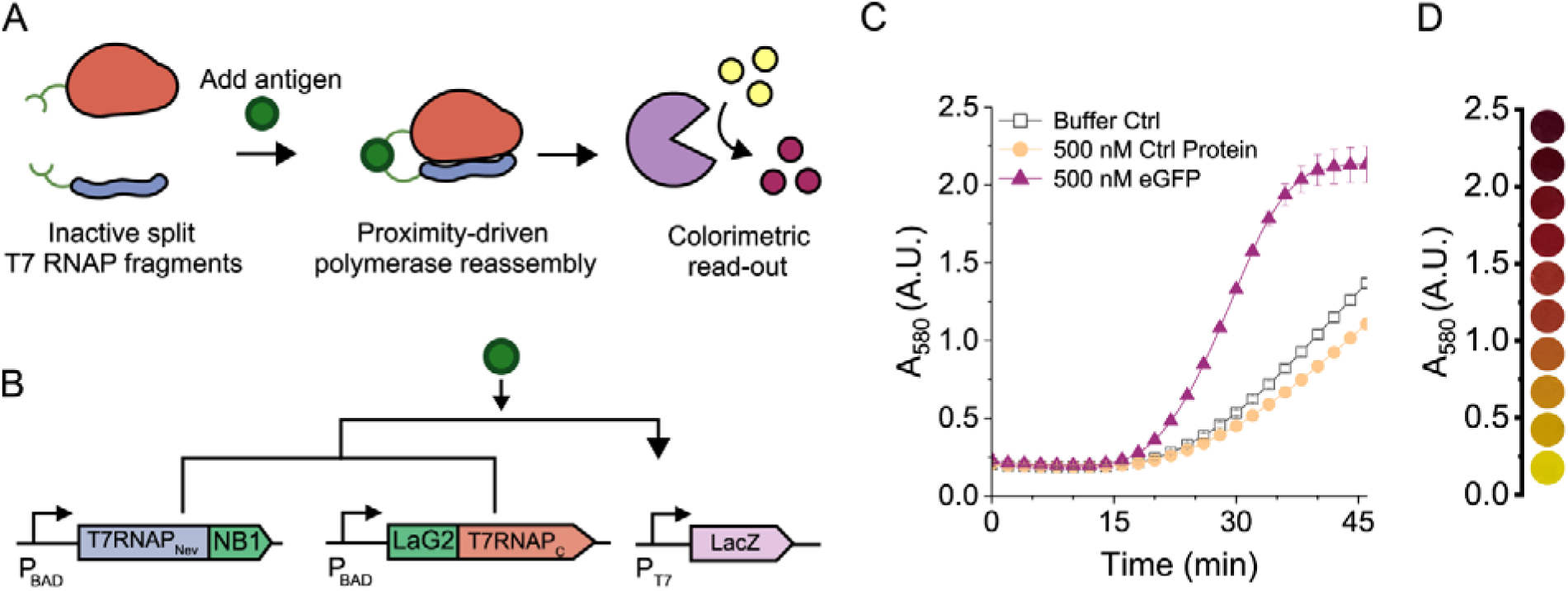
Proof-of-concept eGFP TLISA biosensor. (A) General TLISA schematic. The T7RNAP_Nev_ (blue) fused to a nanobody and the T7RNAP_C_ (red) fused to a separate nanobody should not reassemble in the absence of antigen. In the presence of antigen, both nanobodies bind and bring T7RNAP_Nev_ and T7RNAP_C_ in close proximity, driving reassembly. The reassembled T7RNAP then drives expression of LacZ (purple). LacZ converts the yellow pigment CPRG into purple CPR to generate a range of visually interpretable output colors. (B) Genetic circuit of the proof-of-concept eGFP biosensor. T7RNAP_Nev_ fused to anti-eGFP NB1 and T7RNAP_C_ fused to anti-eGFP LaG2 were expressed from separate plasmids under a native *E. coli* promoter (P_BAD_). In the presence of eGFP, both NB1 and LaG2 bind and bring T7RNAP_Nev_ and T7RNAP_C_ in close proximity, driving reassembly and expression of LacZ off the third, T7-regulated, reporter plasmid. (C) Absorbance readings over time at 37 °C to monitor the conversion of yellow CPRG to purple CPR in the presence of either protein buffer, a control protein (mCherry), or eGFP and 0.1 nM pT7LacZ. Despite some low background reassembly of T7RNAP, target-dependent reassembly and expression is clearly visible. (D) Pictures of visible reaction colors corresponding to different absorbance values. Symbols represent the arithmetic mean ± standard deviation of n=3 technical replicates. Robustness to variation across lysate batches and to reactions on different days is shown in **Fig. S1**.

## Results

### Proof of Principle eGFP TLISA

As a proof-of-principle demonstration, we first aimed to engineer a TLISA biosensor to detect eGFP as a model protein. The TLISA platform was implemented on three plasmids (**Fig. 1B**): one encoding the evolved N-terminal portion (residues 1-179) of T7RNAP (T7RNAP_Nev_) (31) translationally fused to a nanobody (NB), another encoding the C-terminal portion (residues 180-883) of T7RNAP (T7RNAP_C_) translationally fused to a separate NB, and the reporter plasmid encoding β-galactosidase (LacZ) under the regulation of a T7 promoter (pT7LacZ). Anti-eGFP NB sequences were codon optimized for *E. coli* and cloned into plasmids to create a C-terminal fusion on the T7RNAP_Nev_ fragment and an N-terminal fusion on the T7RNAP_C_ fragment. We chose Nb1 (32) and LaG2 (33) as the first NBs to test since their epitopes had little overlap, a feature we predicted to be important to allow both NBs to bind simultaneously. The same 14 amino acid (AA) flexible linker sequence was used on both plasmids to fuse the polymerase to the nanobody. The conversion of the yellow substrate chlorophenol red β-D-galactopyranoside (CPRG) to the purple product chlorophenol red (CPR) was monitored via absorbance at 580 nm to assess sensor functionality. To execute the assay, the T7RNAP fragment fusions were first “pre-expressed” in a CFE reaction step for one hour. Then, reporter plasmid, CPRG, and protein-containing sample were added. Pre-expression of the T7RNAP fragments prior to the final sensing reaction decreases the detection time of the assay. To control for the impacts of the addition of purified target protein and storage buffer on CFE, we used control reactions with either pure buffer or a separate unrelated protein added in place of GFP target.

**Fig. 1C** shows that the addition of eGFP to the TLISA reactions increases the rate of LacZ expression relative to both the buffer and unrelated protein controls, demonstrating TLISA functionality for sensing proteins. The pigmented reporter enables qualitative result interpretation by naked eye observation of reaction color (**Fig. 1D**). We also showed that TLISA reactions are robust to variation across lysate batches and to reactions on different days (**Fig. S1**).

### Flexibility of the TLISA Platform

To demonstrate the modularity of TLISA to work with different nanobodies, new anti-eGFP NB sequences were fused to both the T7RNAP_Nev_ and T7RNAP_C_ fragments and tested for eGFP detection. To facilitate the comparison of the resulting sensors, we used the area between curves (ABC) (**Equation 1**) as a quantitative metric of sensor quality, where ABC ≤ 0 indicates no protein detection and higher ABC indicates more visually distinct responses for a longer period of time—and thus a better sensor. TLISA is rather robust to changes in NB sequence, with several combinations of anti-eGFP NBs producing functional biosensors without any optimization of reaction conditions (i.e., plasmid concentrations and ratios) (**Fig. 2A and Fig. S2**). We note that the T7RNAP_C_ fragments used here contained mutations that propagated during cloning, and these mutations actually improved sensor quality in some instances (**Supplementary Text, Table S1, and Fig. S3**). The best-performing sensor, T7RNAP_Nev_-LaG2/LaG14-T7RNAP_C_, had an ABC of 40.5 and a limit of detection (LOD) of 100 nM eGFP (**Fig. S4**).

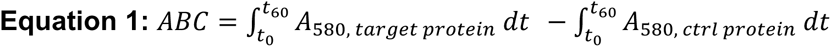

**Fig. 2:**
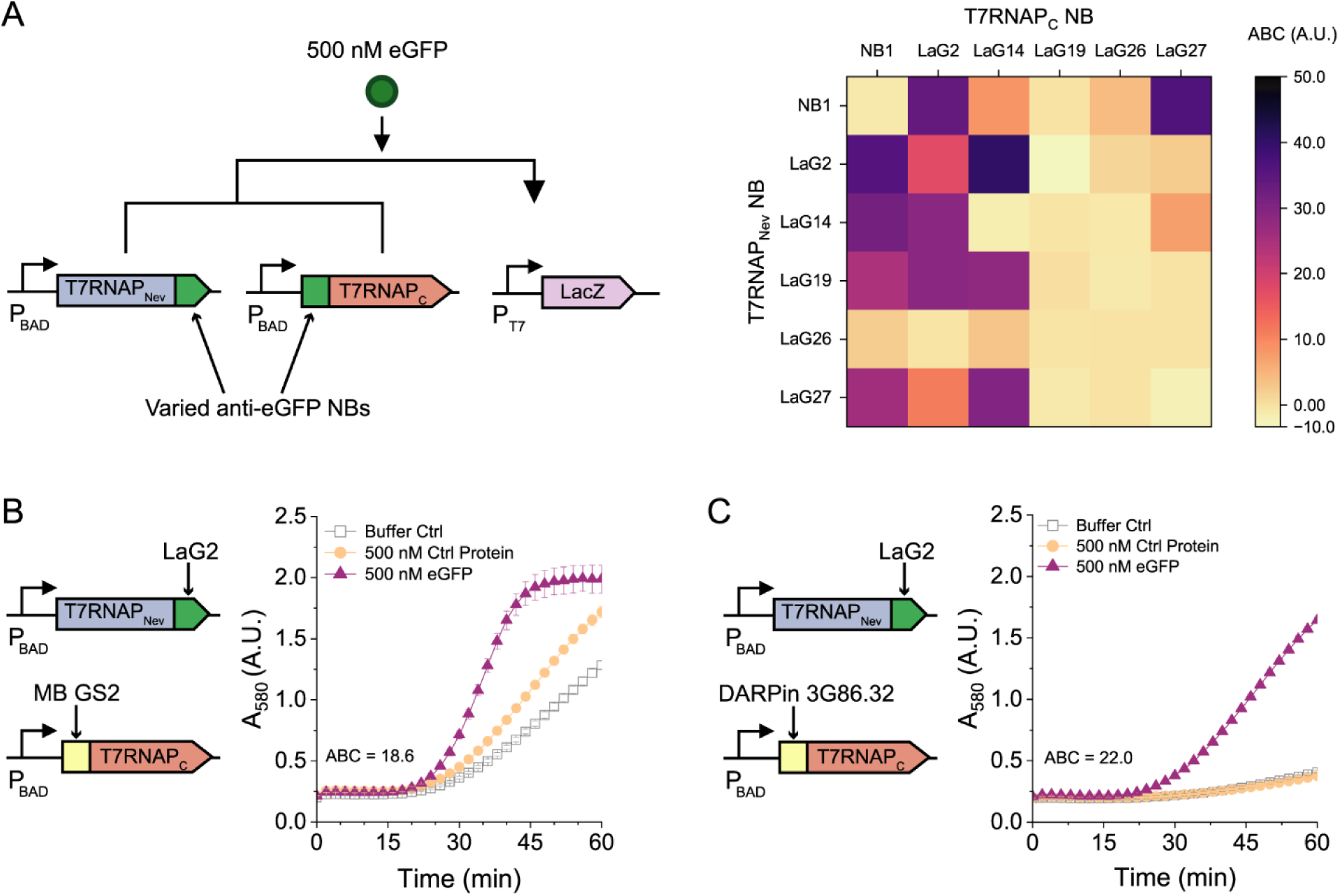
TLISA is robust to different affinity domains. (A) TLISA genetic sensing circuit and ABC (Equation 1) values for all 36 combinations of six anti-eGFP NBs. (B) eGFP TLISA built from a NB (LaG2) and a MB (GS2). (C) eGFP TLISA built from a NB (LaG2) and a DARPin (3G86.32). 500 nM mCherry was used as the control protein in all reactions. All reactions contain 0.1 nM pT7LacZ. Symbols represent the arithmetic mean ± standard deviation of n=3 technical replicates.

These results reveal interesting characteristics of the TLISA platform. First, we expected that to detect monomeric proteins (such as eGFP), the NBs fused to each polymerase fragment would need different epitopes to prevent steric occlusion of one NB by the other, which would otherwise prevent T7RNAP reassembly and reporter expression. The epitopes of the six NBs used here can be organized into three groups based on the general location of their epitopes where LaG14 is in group I; Nb1, LaG19, LaG26, and LaG27 are in group II; and LaG2 is in group III (32,33). The epitopes of LaG19, LaG26, and LaG27 overlap significantly, which could explain why there were no functional sensors created from combinations of these NBs. Surprisingly, a sensor with two LaG2 NBs still yielded significant detection signal. Beyond potential shortcomings in existing epitope maps for these NBs, the mechanism for this signal remains unclear. We also observed that for a given pair of NBs, the ABC could be quite different depending on which NB was on the N versus C fragment. This could be due to differences in protein folding caused by tertiary structure interactions between domains or by improved folding for certain NBs for either N- or C-terminal fusions (as the two fragments had their NBs fused on opposite termini). Lastly, many but not all combinations of NBs result in functional TLISA sensors, and the sensing quality and rate of reaction even for those sensors can vary greatly. Direct eGFP ELISA data confirm that all six anti-eGFP NBs can fold correctly in the CFE environment (**Fig. S5**). These results provide some guidance on which NBs may yield functional TLISA sensors, though it is imperfect. For example, LaG26 had one of the weakest responses in the direct ELISA and also typically had low ABC values in TLISA assays, but NB1 had some of the highest ABC values despite relatively low activity in the direct ELISA. Thus, there are multiple variables that determine whether a NB will prove functional in TLISA, and more exploration is needed to identify characteristics allowing *de novo* prediction of function. Moreover, we note that while all combinations tested in **Fig. 2A** were executed under identical reaction conditions, the performance of individual biosensors can often be further improved simply by optimizing the concentration of plasmid expressing LacZ (**Fig. S6**).

We next showed that TLISA is compatible with different types of protein affinity domains. We implemented two different synthetic protein binding scaffolds as TLISA affinity domains: a monobody and a DARPin (designed ankyrin repeat protein). Monobodies (MBs) are small (ca. 10 kDa) binding proteins based on the fibronectin type III domain. DARPins are based on natural ankyrin repeat protein scaffolds. A new eGFP biosensor using the GS2 MB (34) (**Fig. 2B**) and another sensor using the 3G86.32 DARPin (35) (**Fig. 2C**) were engineered simply by switching them into the T7RNAP_C_ fusion and maintaining all reaction conditions. These results show that TLISA can be robust to different sizes and scaffolds of affinity domains, and that different types of domains can be mixed and matched in one sensor for a plug-and-play platform suitable for rapid re-engineering.

We next investigated the influence of the length of the linker that connects the polymerase fragments to the NBs. Choosing one representative eGFP sensor (T7RNAP_Nev_-NB1/LaG2-T7RNAP_C_), we altered linker lengths from 14 AAs to either 7 or 28 AAs for each polymerase fragment and tested all possible combinations for eGFP detection. Linker length affected detection functionality differently for the two polymerase fragments (**Fig. 3A**). Using a 28 AA linker on the T7RNAP_Nev_ fragment abrogated detection in all combinations, while longer linkers on the T7RNAP_C_ fragment generally yielded strong detection. The best combination used a 7 AA linker on the T7RNAP_Nev_ fragment and a 28 AA linker on the T7RNAP_C_ fragment (**Fig. 3B**). Again, we suspect that these observations could be due to differences in protein folding and stability for the fused constructs. While these results may not generalize to all combinations of NBs and their respective epitope locations, they do indicate that linker length could be used as a tuning parameter to improve signal-to-noise for the same pair of NBs.

**Fig. 3:**
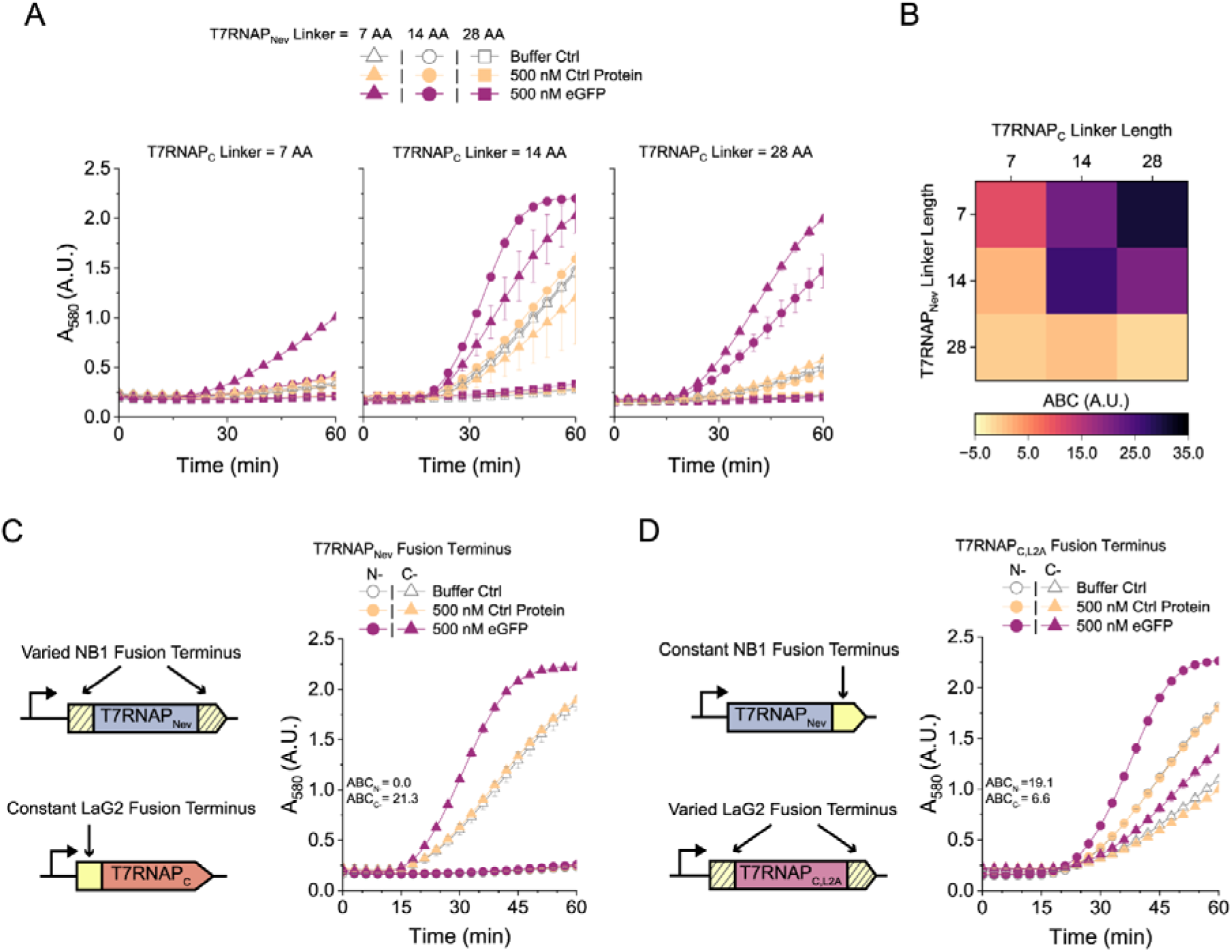
Impacts of linker length and fusion termini on sensor performance. (A) Absorbance data of T7RNAP_Nev_-NB1/LaG2-T7RNAP_C_ TLISAs with different linker lengths. Different shapes represent different linker lengths for T7RNAP_Nev_-NB1. Each graph represents a different linker length for LaG2-T7RNAP_C_. (B) ABC values for the data presented in (A). (C) Genetic circuits indicating the varying locations of the NB on the T7RNAP_Nev_ fragment and absorbance data for TLISA reactions with NB1 fused to either the N- (circles) or C-terminus (triangles) of T7RNAP_Nev_ used with LaG2-T7RNAP_C_. ABC values for the two sensors are displayed in the graph. (D) Genetic circuits indicating the varying locations of the NB on the T7RNAP_C,L2A_ fragment and absorbance data for TLISA reactions with LaG2 fused to either the N- (circles) or C-terminus (triangles) of T7RNAP_C,L2A_ used with T7RNAP_Nev_-LaG19. ABC values for the two sensors are displayed in the graph. All reactions contained 0.1 nM pT7LacZ except for the C-terminal NB fusion data in D, which used 0.2 nM pT7LacZ. 500 nM mCherry was used as the control protein for all reactions. Symbols represent the arithmetic mean ± standard deviation of n=3 technical replicates.

We also investigated the impacts of changing the terminus for the NB fusion for each polymerase fragment. All experiments thus far used the C-terminus of T7RNAP_Nev_ and N-terminus of T7RNAP_C_ for fusions, based on previous reports for *in vivo* split T7RNAP biosensors (30,31). However, as we have already shown, functionality is dependent on the local fusion context, so changing fusion locations could potentially yield even better sensors. We tested all four combinations of fusion termini with the split T7RNAP system using the wt version of both fragments; this wt version of the T7RNAP_N_ fragment has not been evolved to minimize spontaneous reassembly, and thus provides a way to assess whether functional polymerase can reassemble at all in a CFE system. We found that both N- and C-terminal fusions to wt T7RNAP_N_ yield reassembly of functional polymerase, while T7RNAP_C_ does not tolerate C-terminal fusions (**Fig. S7**), consistent with previous reports (36). We then also tested a second, mutated T7RNAP_C_ fragment (T7RNAP_C,L2A_) whose mutations enable C-terminal fusions (36); we validated spontaneous reassembly in a CFE system with a NB fused to the C-terminus of the T7RNAP_C,L2A_ fragment (**Fig. S7**) even though this was not possible with the original T7RNAP_C_ fragment.

Assessment of TLISA functionality for NB fusions on different termini for T7RNAP_Nev_ and T7RNAP_C,L2A_ showed some degree of robustness to these changes. Although the wt T7RNAP_N_ tolerated both N- and C-terminal fusions, we observed no detection and minimal baseline activity when fusing NBs to the N-terminus of T7RNAP_Nev_ (**Fig. 3C**). This loss of activity may be due to protein folding differences between the wt T7RNAP_N_ and T7RNAP_Nev_. On the other hand, we demonstrated functional eGFP biosensors with NB fusions to both the N- and C-terminus of T7RNAP_C,L2A_ (**Fig. 3D**). An N-terminal NB fusion on T7RNAP_C,L2A_ resulted in a functional eGFP sensor using the established reaction conditions without any optimization while the pT7LacZ reporter plasmid concentration needed to be increased to observe eGFP detection when using the C-terminal NB fusion on the T7RNAP_C,2LA_. Thus, future TLISA designs could use selection of termini for fusions as another possible avenue for optimization.

### TLISA can easily be reengineered to detect different antigens

We next showed that the TLISA platform can be used to detect a different protein antigen. The modular design nature of TLISA makes this simple: we cloned new plasmids with anti-mCherry NB fusions to the polymerase fragments to create a new TLISA sensor for mCherry (**Fig. 4A**). The epitopes of these NBs were not characterized at the time of selection, so of the available NBs we choose the four with the lowest K_d_ values (33) and validated proper CFE with an ELISA (**Fig. S8**). We tested for mCherry detection using all 16 possible nanobody combinations. The best-performing sensor, T7RNAP_Nev_-LaM4/LaM2-T7RNAP_C_, had an ABC of 54.9 (**Fig. 4A**) and an LOD of 50 nM mCherry (**Fig. S9**). Similar to eGFP sensors, many different combinations of anti-mCherry nanobodies can be used to produce functional mCherry sensors (**Fig. 4B and Fig. S10**). The epitopes of LaM2, LaM4, and LaM6 were later reported, revealing that the epitopes of LaM2 and LaM6 overlap significantly while the epitope of LaM4 is separate (37) and thus explaining why sensors combining LaM2 and LaM6 yielded no detection. We also showed that TLISA can use reporter proteins other than LacZ, implementing superfolder GFP (sfGFP) as a reporter protein for an mCherry TLISA now that eGFP was no longer the target antigen (**Fig. S11**).

**Fig. 4:**
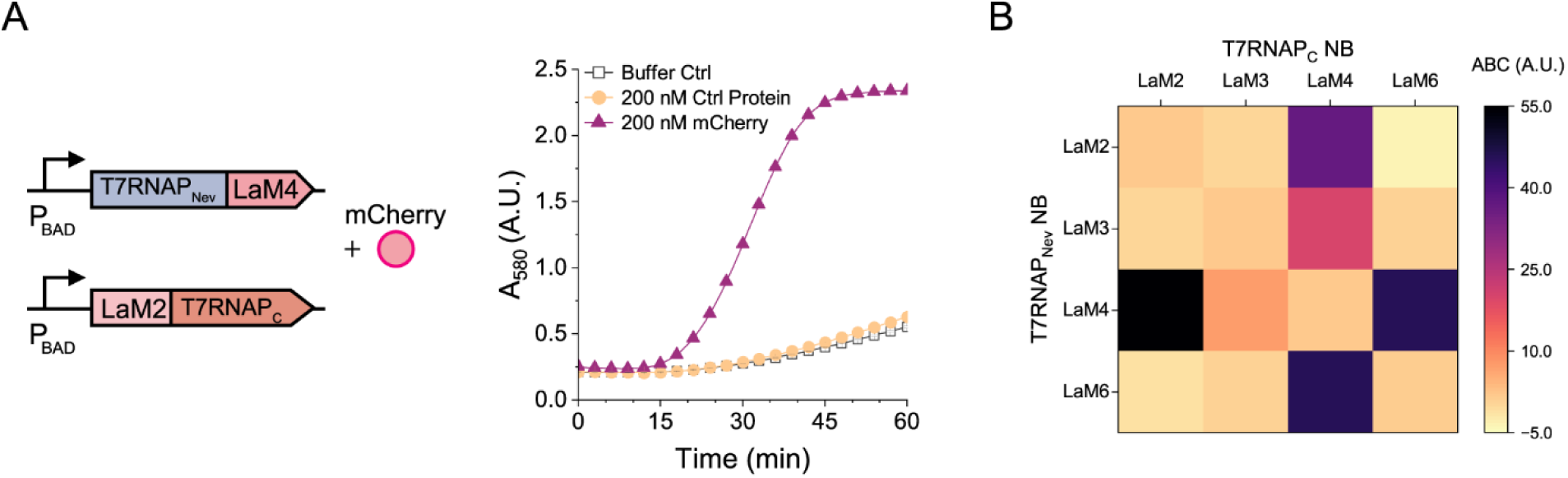
TLISA is modular for detection of different protein targets. (A) Genetic circuit for sensing mCherry and corresponding absorbance versus time data. (B) ABC values for all combinations of mCherry NBs. All reactions had 0.05 nM pT7LacZ. 200 nM eGFP was used as the control protein for all reactions. Symbols represent the arithmetic mean ± standard deviation of n=3 technical replicates.

### Rapid development of clinically relevant biosensors

Next, we showed that it is straightforward to apply the TLISA platform for detection of clinically relevant targets. We first engineered a biosensor to detect the receptor binding domain (RBD) of the severe acute respiratory syndrome coronavirus 2 (SARS-CoV-2) spike (S) protein. NBs VHHE and VHHV were chosen due to their high binding affinities and non-overlapping epitopes (38). NB sequences were codon optimized, cloned into T7RNAP fragment plasmids as translational fusions, and tested for the ability to detect soluble RBD. By fusing VHHE to T7RNAP_Nev_ and VHHV to T7RNAP_C_, we successfully engineered a TLISA RBD biosensor and observed maximum signal-to-noise in 35 minutes (**Fig. 5A**). We also observed significant RBD detection when exchanging the fragments to which the NBs were fused (i.e., fusing VHHV to T7RNAP_Nev_ and VHHE to T7RNAP_C_), though the signal was not as strong (**Fig. S12**). Since we previously showed that DARPins can be efficiently incorporated into the TLISA platform (**Fig. 2C**), we selected an existing DARPin (FSR22) (39) against the RBD and fused it to T7RNAP_Nev_ to generate another functional TLISA sensor, which had lower leak than the dual-NB version (**Fig. 5B**). Its LOD was 200 nM RBD (**Fig. S13**); while this is greater than the LOD of commercialized rapid antigen tests (40) and would thus require further engineering for clinical deployment, we have nonetheless shown that TLISA enables quick and easy creation of biosensors against targets of clinical interest without extensive optimization. Additionally, **Fig. 5A** and **Fig. 5B** show that both RBD biosensors are robust to day-to-day experimental variation.

**Fig. 5:**
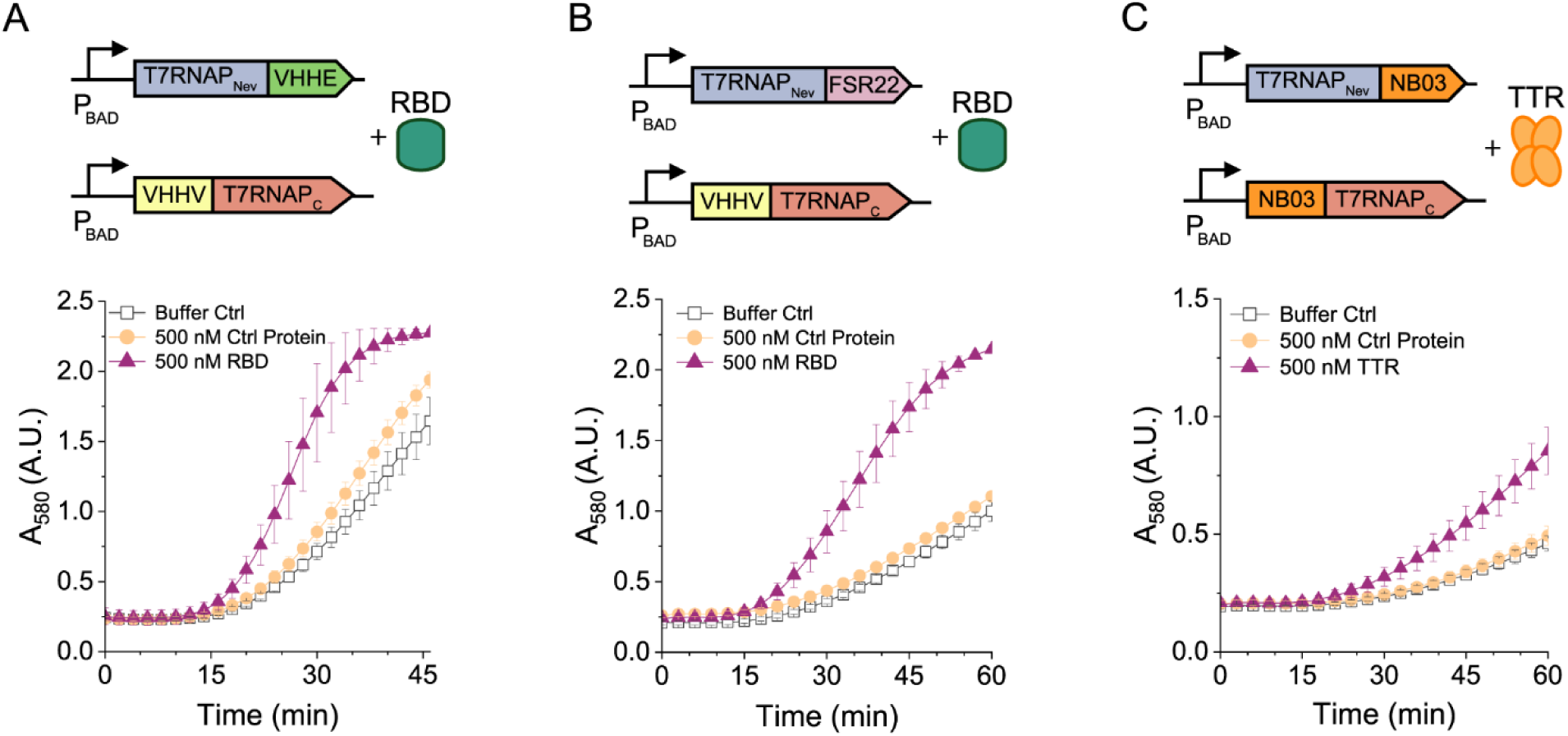
TLISA can be easily re-engineered to create biosensors for clinically relevant protein biomarkers. (A) Genetic circuit and absorbance versus time of a TLISA sensor to detect SARS-CoV-2 RBD using two NBs (VHHE and VHHV) or (B) one NB and one DARPin (FSR22). (C) Genetic circuit and absorbance versus time data of a TLISA sensor detecting transthyretin (TTR) using NB03 on both polymerase fragments. Both SARS-CoV-2 biosensors used 0.1 nM pT7LacZ. The TTR biosensor used 0.12 nM pT7LacZ. 500 nM mCherry was used as the control protein in all reactions. Symbols represent the arithmetic mean ± standard deviation of n=6 replicates performed on different days.

We then created a biosensor for another clinically relevant target, transthyretin (TTR). Serum TTR has historically been used as a biomarker to determine overall nutritional status (41). Malnutrition is the direct cause of about 300,000 deaths per year (42,43), primarily in the developing world. Because TTR is a homotetramer, we posited that TLISA could work even with the same NB fused to both plasmids, simplifying sensor design. We thus chose NB03 (44) as our protein affinity domain for both T7RNAP fragments. The first TTR sensor we constructed successfully detected TTR within 1 hour with robustness to day-to-day variation (**Fig. 5C**). The signal-to-noise ratio was relatively low, though, so we attempted some basic sensor optimization. We decreased the linker length from 14 AA to 7 AA on the T7RNAP_Nev_-NB03 fragment based on our eGFP sensor results (**Fig. 3B**). This resulted in a much faster reaction time, and the ABC increased from 6.7934 to 16.7577 (**Fig. S14**). Decreasing the concentration of the pT7LacZ reporter plasmid from 0.12 nM to 0.05 nM to slow the reaction rate yielded a similar ABC but with less leak and thus with potentially improved visual interpretability (**Fig. S14**). This TLISA biosensor has an LOD of 500 nM TTR, which is well below the healthy range of serum TTR concentration (3 μM to 8 μM) (45).

### Towards point-of-care use

TLISA is also robust for detection in complex biological sample matrices that would be used in a clinical diagnostic device. The established reaction protocol remains the same, but the protein to be detected is spiked into a biological sample matrix that is added after the pre-expression (**Fig. 6A**). For field deployment, pre-expression would be a part of the manufacturing process, not an end user step. Lysates enriched with NB-T7RNAP fragments can be used to eliminate the need for pre-expression, but pre-expression was used here to characterize many different sensors more efficiently. To account for increased RNAse activity in human serum and saliva, we added 1% Murine RNAse inhibitor per previously established protocols (8). For an mCherry sensor, we added either 10% or 20% pooled human saliva or pooled human serum spiked with target protein.

**Fig. 6:**
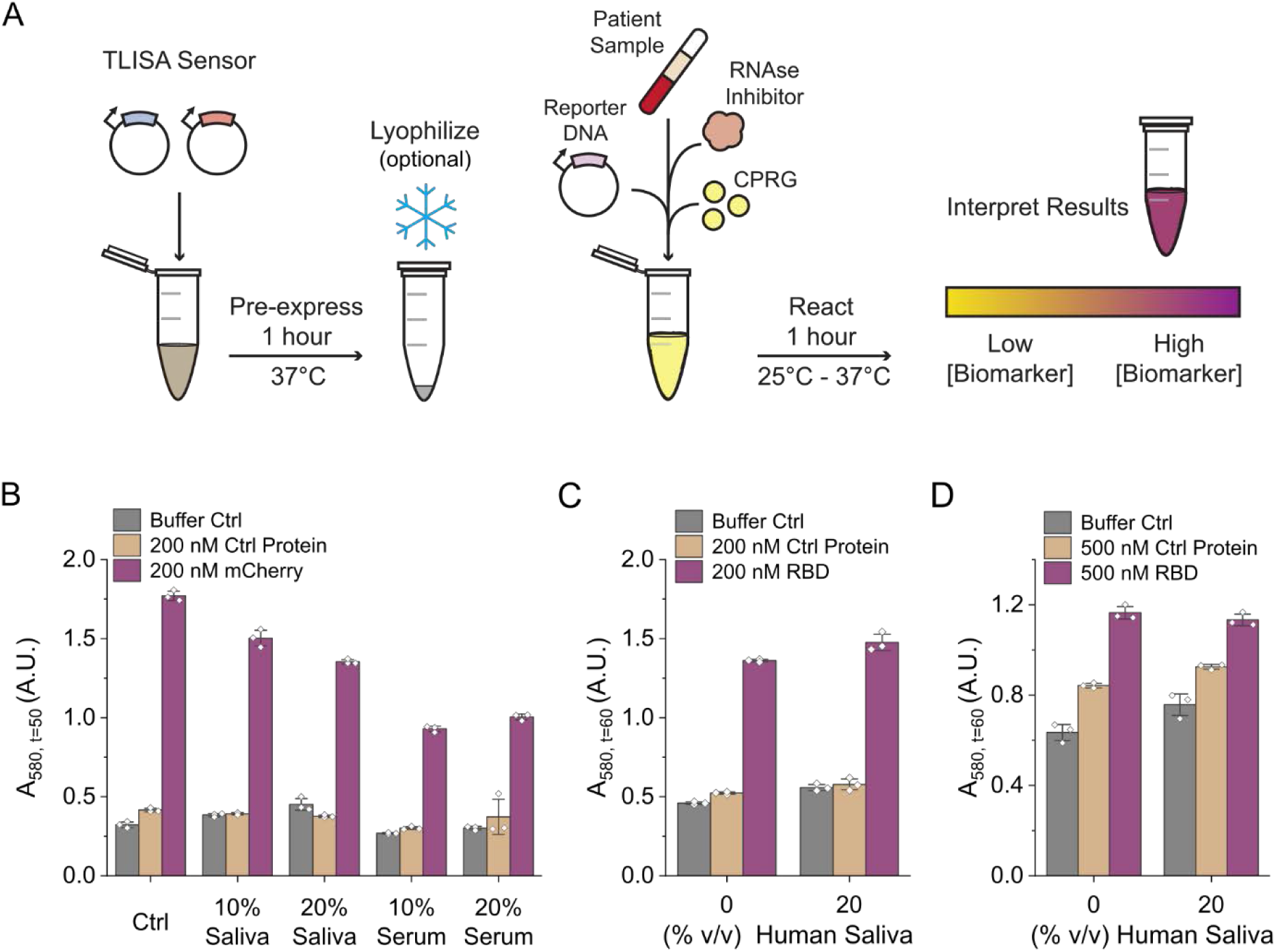
TLISA functions robustly in complex sample matrices and after lyophilization. (A) Schematic of TLISA protocol when sensing proteins in biological samples. (B) End-point absorbance values after 50 minutes for an mCherry TLISA in various concentrations of either pooled human serum or saliva with 1% v/v RNAse inhibitor. (C) Absorbance values after 60 minutes in a SARS-CoV-2 RBD TLISA in either 0% or 20% v/v pooled human saliva with 1% v/v RNAse Inhibitor. (D) Absorbance values of a lyophilized SARS-CoV-2 RBD TLISA reaction 60 minutes after rehydration with either 0% or 20% v/v pooled human serum, 0.5% v/v RNAse Inhibitor, and 0.75 nM pT7LacZ. 200 nM eGFP was used as a control in (B). 200 nM mCherry was used as a control protein for (C) and (D). Bars represent the arithmetic mean ± standard deviation of n=3 technical replicates (white diamonds).

**Fig. 6B** shows successful detection of mCherry in all conditions. The substantially lower final absorbance values observed when running reactions in serum vs. defined aqueous mixtures are likely due to the binding of serum albumin to CPR eliciting a shift in the absorption spectrum. We have previously shown that the addition of naproxen to reactions with CPRG in serum restores proper color (9).

Moreover, TLISA can detect clinically relevant human biomarkers in their corresponding complex sample matrices. To demonstrate this, we validated function of a sensor targeting SARS-CoV-2 RBD (T7RNAP_Nev_-FSR22/VHHV-T7RNAP_C_) after spiking in 20% human saliva. The addition of 20% human saliva did not impact RBD detection (**Fig. 6C**). Towards our field-friendly vision, we also confirmed that TLISA can detect 200 nM RBD within a 20% human saliva matrix when reacted at room temperature (25 °C) (**Fig. S15**).

Finally, we demonstrated that TLISA has the potential for use in minimally equipped settings by showing it retains function after lyophilization. Lyophilization of cell-free reactions helps enable stable room temperature storage of biosensors that can then be rehydrated and used at the POC. We lyophilized our TLISA reactions after the pre-expression step (**Fig. 6A**), just before sample would be added. We first confirmed that pre-expressed T7RNAP and split T7RNAP polymerase fragments can be lyophilized after pre-expression and properly rehydrated (**Fig. S16A and S16B**). Then, we showed that TLISA sensing reactions are robust to lyophilization; we demonstrated this with the T7RNAP_Nev_-LaM4/LaM2-T7RNAP_C_ mCherry sensor (**Fig. S16C**) as well as the SARS-CoV-2 RBD sensor (**Fig. S16D**). Finally, after adjusting RNAse inhibitor and pT7LacZ concentrations, we demonstrated detection of SARS-CoV-2 RBD in 20% human saliva from a lyophilized TLISA reaction (**Fig. 6D**). Taken together, rapid detection of human biomarkers in biological matrices from lyophilized reactions demonstrates that TLISA is well-suited for future clinical diagnostic applications.

## Discussion

We have developed and characterized a novel, highly adaptable and generalizable CFE biosensing platform for protein detection. TLISA can detect different antigens using numerous combinations of nanobodies, monobodies, and DARPins, and can be linked to different reporter proteins, demonstrating the truly modular nature of this platform. Importantly, TLISA is robust to both day-to-day experimental variation and variation between different batches of in-house prepared crude lysate with only minor changes in sensor performance. It also functions robustly at room temperature, in complex biological matrices, and after lyophilization, achieving room temperature colorimetric detection of SARS-CoV-2 RBD in human saliva in just one hour.

While TLISA clearly shows promise as an impactful POC diagnostic tool, performance improvements will be critical for future in-field deployment. Although many of the sensors presented here exhibited low leak and high signal in the presence of target antigen, some had less desirable signal-to-noise ratios. Using different NB fusions, linker lengths, and reporter plasmid concentrations can help tune performance. High-throughput studies to identify trends between NB sequence and signal-to-noise ratio could provide particularly useful insights useful for *de novo* engineering of sensors for new targets. New strategies to decrease leak due to the spontaneous RNAP reassembly would also significantly improve sensitivity and usability in some applications. However, there are numerous applications where sensitivity at levels similar to the TLISA examples presented here would be effective, like for the detection of the inflammation biomarker C-reactive protein, whose elevated levels are defined as above 5 mg/L (corresponding to 40 nM) (46). Additionally, the current linear range of detection is narrower than that of an ELISA. Broadening this range would increase the potential application space of TLISA. Nonetheless, the dose responses shown here provide an important feature that standard lateral flow assays for POC protein detection lack.

Even beyond POC applications, though, TLISA could provide unique capabilities in clinical laboratories as well. Compared to ELISA assays that are widely used for measuring proteins, the TLISA workflow is much simpler and much less subject to inter-operator variability, which would improve reliability of results. In addition, at 10 μL the volume necessary for a TLISA assay is extremely small, which could provide significant advantages for a clinical assay as more measurements could be taken from the same sample. This benefit would be particularly impactful for neonatal care where daily blood draw volumes can be limited to just a few mL.

The initial characterization presented here has only begun to explore the potential breadth of this platform. The successful use of nanobodies, DARPins, and monobodies in TLISA suggests a vast repertoire of potential affinity domain fusions. We anticipate that many other scaffolds such as single-chain variable fragments (scFvs), affibodies, and anticalins can also be used to develop novel protein sensors. Furthermore, TLISA can potentially be used to engineer protein-mediated OR gates by using bispecific affinity domains, as well as AND gates by splitting the T7RNAP into additional fragments at previously identified cut sites (47). Finally, while we focused our efforts on sensing proteins so as to fill a significant gap in the field, using affinity domains against non-protein antigens (e.g., small molecules, glycans, etc.) could expand TLISA to sense a variety of diverse analytes and give it an even broader potential impact.

## Materials and Methods

### Bacterial Strains and Plasmid Preparation

DNA oligonucleotides for cloning and sequencing were synthesized by Eurofins Genomics. T7RNAP fragments were amplified via PCR from the BL21 (DE3) genome. Gene strands for T7RNAPNev, and all nanobody, monobody, and DARPin sequences were codon optimized and synthesized by Twist Biosciences. Plasmids were cloned by either Gibson Assembly (48) or blunt-ended ligation using the pJL1 plasmid backbone. *E. coli* strains DH10β and DH5α were used for cloning and plasmid preparations. Isolated colonies were grown overnight in LB medium with either kanamycin sulfate (33 μg/mL) or tetracycline (15 μg/mL). Plasmid DNA from overnight cultures was purified using EZNA mini prep columns (OMEGA Bio-Tek). Plasmid sequences were verified with Sanger DNA sequencing (Eurofins Genomics). Sequence-confirmed plasmids were then purified using EZNA midiprep or maxiprep columns (OMEGA Bio-Tek), followed by isopropanol and ethanol precipitation. The purified DNA pellet was reconstituted in elution buffer, measured on a Nanodrop 2000 for concentration, and stored at −20 °C until use. *E. coli* strain BL21 ΔlacIZYA was created by lambda red recombination (49) and used for in-house cell-free lysate preparation.

### Preparation of cell-free lysate

Bacterial lysate for all experiments was prepared as described by Kwon and Jewett (50) with a few protocol modifications. BL21 ΔlacIZYA cells were grown in 2xYTP medium at 37 °C and 220 rpm to an optical density (OD) between 1.5-2.0, corresponding to the mid-exponential growth phase. Cells were centrifuged at 4 °C and 2700 x *g* and washed via resuspension with S30 buffer (10 mM tris-acetate [pH 8.2], 14 mM magnesium acetate, 60 mM potassium acetate, and 2 mM dithiothreitol). These centrifugation and wash steps were repeated twice for a total of three S30 washes. After the final centrifugation, the wet cell mass was measured, and cells were resuspended in 1 mL S30 buffer per 1 g of wet cell mass. The cellular resuspension was divided into 0.5 mL aliquots. Cells were lysed using a Q125 sonicator (Qsonica) at a frequency of 20 kHz and 50% amplitude. Cells were sonicated on ice with cycles of 10 seconds on and 10 seconds off, delivering approximately 150 J, at which point the cells appeared visibly lysed. An additional 4 mM dithiothreitol was added to each tube immediately after lysing, and the sonicated mixture was then centrifuged at 12,000 x *g* and 4 °C for 10 minutes. After centrifugation, the supernatant was divided into 1 mL aliquots, and incubated at 37 °C and 220 rpm for 80 minutes. After this runoff reaction, the lysate was centrifuged at 12,000 x *g* and 4 °C for 10 minutes. The supernatant was removed and loaded into a 10 kDa molecular weight cutoff dialysis cassette (Thermo Fisher). Lysate was dialyzed in 1 L of S30B buffer (14 mM magnesium glutamate, 60 mM potassium glutamate, 1 mM dithiothreitol, and pH-corrected to 8.2 with tris) at 4 °C for 3 hours. Dialyzed lysate was removed and centrifuged at 12,000 x *g* and 4 °C for 10 minutes. The supernatant was removed, aliquoted, and stored at −80 °C until use.

### TLISA Cell-free Reactions

All TLISA reactions consisted of a pre-expression reaction and a final sensing reaction. For the pre-expression, cell-free reactions were assembled in 1.5 mL centrifuge tubes as previously described by Kwon and Jewett (50) with 10 mM arabinose and equimolar concentrations (25 nM) of T7RNAPNev-NB and NB-T7RNAPC plasmids. Reactions were pre-expressed at 37 °C for one hour. Then, pT7LacZ, 0.6 mg/mL CPRG, and purified protein were added to each tube for the final sensing reaction. Each cell-free reaction had a final volume of 10 uL and was pipetted into a clear-bottomed 384-well plate. The final sensing reaction was carried out in the BioTek plate reader at 37 °C for one hour with absorbance measurements taken every minute. Plates were sealed with a transparent adhesive film to prevent evaporation. For reactions containing biological samples, the final sensing reaction contained either 10% or 20% pooled human serum (MP Biomedicals) or 10% or 20% pooled human saliva (Innovative Research Inc.) and RNAse inhibitor, murine (New England BioLabs).

### Protein Purification

For the expression and purification of RBD, a plasmid containing amino acids 319-541 of the SARS-CoV-2 S protein (UniProt P0DTC2) followed by a GGSGG linker, a SpyTag, and a 6xHis Tag was transfected into Expi293F suspension cells with the ExpiFectamine 293 transfection kit (Gibco) according to the manufacturer’s protocol. This plasmid was codon optimized for expression in mammalian cells and synthesized by Gene Universal Inc. (Newark, DE). Five days after transfection, the culture was centrifuged for 5 minutes at 5000 x *g*, and the supernatant was thoroughly dialyzed into PBS. Ni-NTA resin was equilibrated with 10 column volumes (CVs) of IMAC binding buffer (150 mM Tris, 150 mM NaCl, 20 mM imidazole, pH 8.0), then dialyzed supernatant was added to the resin. The resin was washed with 20 CVs of binding buffer, and the protein was eluted with 10 CVs of elution buffer (150 mM Tris, 150 mM NaCl, 400 mM imidazole, pH 8.0). Eluted protein was concentrated in a 10 kDa MWCO Amicon spin filter (EMD Millipore) to < 1 mL. Concentrated RBD was injected onto a Superdex Increase 200 10/300 GL (Cytiva) size exclusion column to remove any remaining impurities. Concentration was measured by bicinchoninic acid (BCA) assay, and purity was assessed by SDS-PAGE (**Fig. S17**).

For the expression and purification of eGFP, mCherry, and TTR, plasmids expressing 6xhis-tagged proteins under pBAD regulation were transformed into BL21 ΔlacIZYA cells. Single colonies were then grown overnight in 50 mL of 2xYTP medium and 15 μg/mL tetracycline at 37 °C and 220 rpm. The next day, 5 mL of the overnight culture was diluted in 500 mL 2xYTP medium with tetracycline and incubated at 37 °C and 220 rpm in a 1 L baffled flask. Once the OD reached 0.4-0.6, 2 mM arabinose was added to the culture to induce protein expression. Flasks were then transferred to a separate incubator and grown overnight at 25 °C and 180 rpm. The next day, cultures were transferred to 50 mL tubes and centrifuged at 2700 x *g* and 4 °C for 15 minutes. Cells were washed in 1xPBS and centrifuged again under the same conditions. The supernatant was discarded, and cells were resuspended in 2 mL of lysis buffer (50 mM disodium phosphate, 500 mM sodium chloride, 10 mM imidazole, pH 8) per 1 g of wet cell mass. Cells were lysed via sonication using the conditions specified for cell-free lysate preparation except without the addition of dithiothreitol. Lysed cells were centrifuged at 12,000 x *g* and 4 °C for 15 minutes. Lysates were loaded onto a pre-equilibrated Ni-NTA column and washed with 20 CVs of binding buffer. Each protein was eluted with 10 CVs of elution buffer, then concentrated by spin filter to < 1 mL. Proteins were further purified by size exclusion chromatography. Fractions containing high concentrations of protein were collected and combined. Protein purity was assessed via SDS-PAGE (**Fig. S17**). Protein concentration was determined by absorbance at 280 nm.

### Lyophilization

TLISA pre-expression reactions were prepared as previously described in PCR tubes. After one hour of pre-expression, tubes were transferred to a prechilled Labconco Fast-freeze flask and stored at −80 °C until frozen. The flask was then connected to a Labconco FreeZone benchtop freeze dryer and samples lyophilized at −50 °C and 0.02 mbar for 3 hours. Samples were then removed and rehydrated on ice for the final reaction.

### ELISAs

NB_LacZ fusion proteins were expressed in cell-free reactions. Cell-free reactions were assembled in 1.5 mL centrifuge tubes as previously described by Kwon and Jewett(50) with 5 nM of the NB-LacZ fusion plasmid. Reactions were incubated at 30 °C overnight. Then, cell-free reactions were centrifuged at 12,000 x *g* and 4 °C for 15 minutes. The supernatant was diluted 1:4 in 1x PBS. 100 uL of 1 mg/mL of purified 6xhis-tagged proteins were added to a Ni-coated 96-well plate and incubated for 2 hours at room temperature on a rocker. Wells were washed four times with wash buffer (1xPBS with 0.05% Tween-20) to remove unbound protein. 100 uL of the diluted NB_LacZ was added to each well and incubated for two hours at room temperature on a rocker. Wells were washed four times with wash buffer to remove unbound NBs. 100 uL of CPRG was added to each well. The plate was incubated for a final time at 37 °C without shaking. End absorbance was measured using the BioTek plate reader.

### Color images and processing

Pictures of colorimetric reactions were taken with an iPhone 7. Pictures represent 10 μL reactions in clear, flat-bottomed 384-well plates. The centers of the wells of interest were cropped using Adobe Illustrator and combined to make color arrays. A brightness filter was uniformly applied to photos to make them better resemble actual appearance.

## Acknowledgments

The authors thank Michael C. Jewett and Julius B. Lucks for their gift of the pJL1 plasmid. We also thank Bryan C. Dickinson for his gift of the plasmid containing T7RNAP_C,L2A_.

## Author contributions

Conceptualization: MAM, MPM, MPS

Investigation: MAM, RCLL

Resources: MAM, ATP, KL

Formal Analysis: MAM Writing—original draft: MAM

Writing—review & editing: MAM, ATP, KL, MPM, RSK, MPS

Visualization: MAM

Supervision: MPS

Funding acquisition: RSK, MPS

## Data and materials availability

All data are available in the main text or the supplementary materials.

