## Supplementary Material for "A modular cell-free protein biosensor platform using split T7 RNA polymerase"

Megan A. McSweeney *et al*.

Corresponding

**This PDF file includes:**

Supplementary Text
Figs. S1 to S17
Tables S1 to S5

**Supplementary Text**

During this study, and after most of the data were collected, we noticed mutations in the gene for T7RNAP$_C$ on several plasmids. For full transparency, we have included **Table S1** that indicates which mutations were present on each plasmid used to generate the data in every figure. None of the plasmids used in this study have mutations in the NB, MB, or DARPin sequences. Additionally, there were no mutations in any plasmids encoding for the T7RNAP$_N$ or T7RNAP$_{Nev}$ fragments. Plasmids not listed in this table did not have any mutations.

After identifying mutations in these plasmids, we corrected the mutations on three NB-T7RNAP$_C$ plasmids to test how these mutations impact TLISA functionality. Correcting the mutations on the LaG2-T7RNAP$_C$ plasmid resulted in a dysfunctional sensor (**Fig. S3A**). We decreased the reporter plasmid concentration to reduce leak but still did not observe any eGFP detection under any conditions (**Fig. S3B**). For the eGFP sensor that uses a MB fusion to the T7RNAP$_C$ (T7RNAP$_{Nev}$-LaG2/GS2-T7RNAP$_C$), correcting the plasmid mutations improved the rate of reaction, and was still able to detect 500 nM eGFP (**Fig. S3C**). The eGFP sensor that uses a DARPin fusion to the T7RNAPC (T7RNAP$_{Nev}$-LaG2/3G86.32-T7RNAP$_C$) also failed to detect eGFP after the mutations were corrected (**Fig. S3D**).

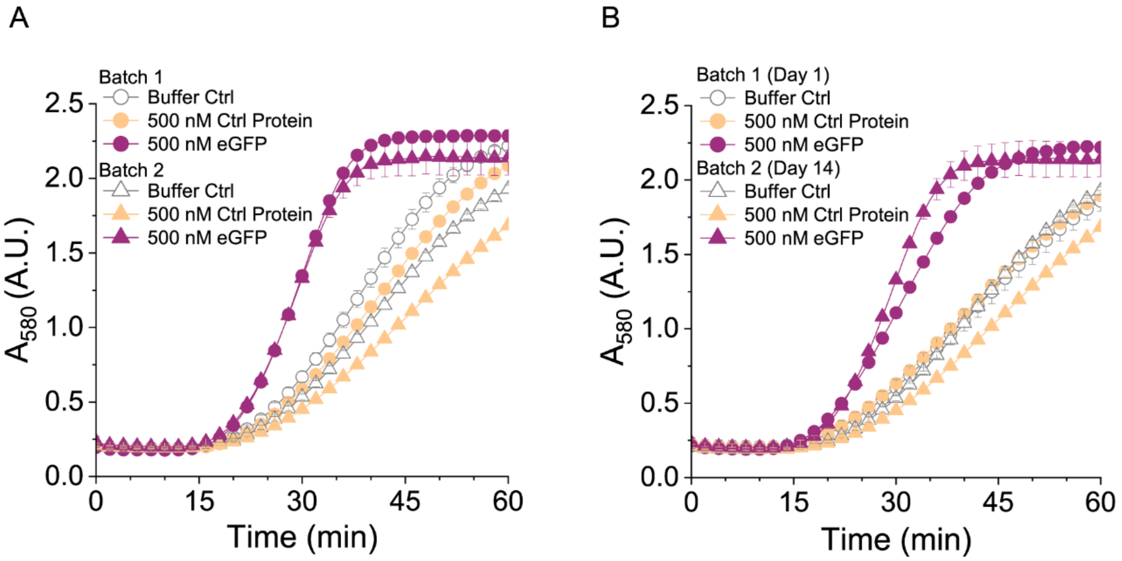


**Fig. S1.**

TLISA performance has minimal variation across different batches of crude lysate and different reaction days. (A) Absorbance data for a T7RNAP$_{Nev}$-NB1/LaG2-T7RNAP$_{C}$ TLISA using two different batches of in-house prepared crude lysate. (B) Absorbance data for a T7RNAP$_{Nev}$-NB1/LaG2-T7RNAP$_{C}$ TLISA using two different batches of in-house prepared crude lysate performed two weeks apart. Symbols represent the arithmetic mean ± standard deviation of n=3 technical replicates.

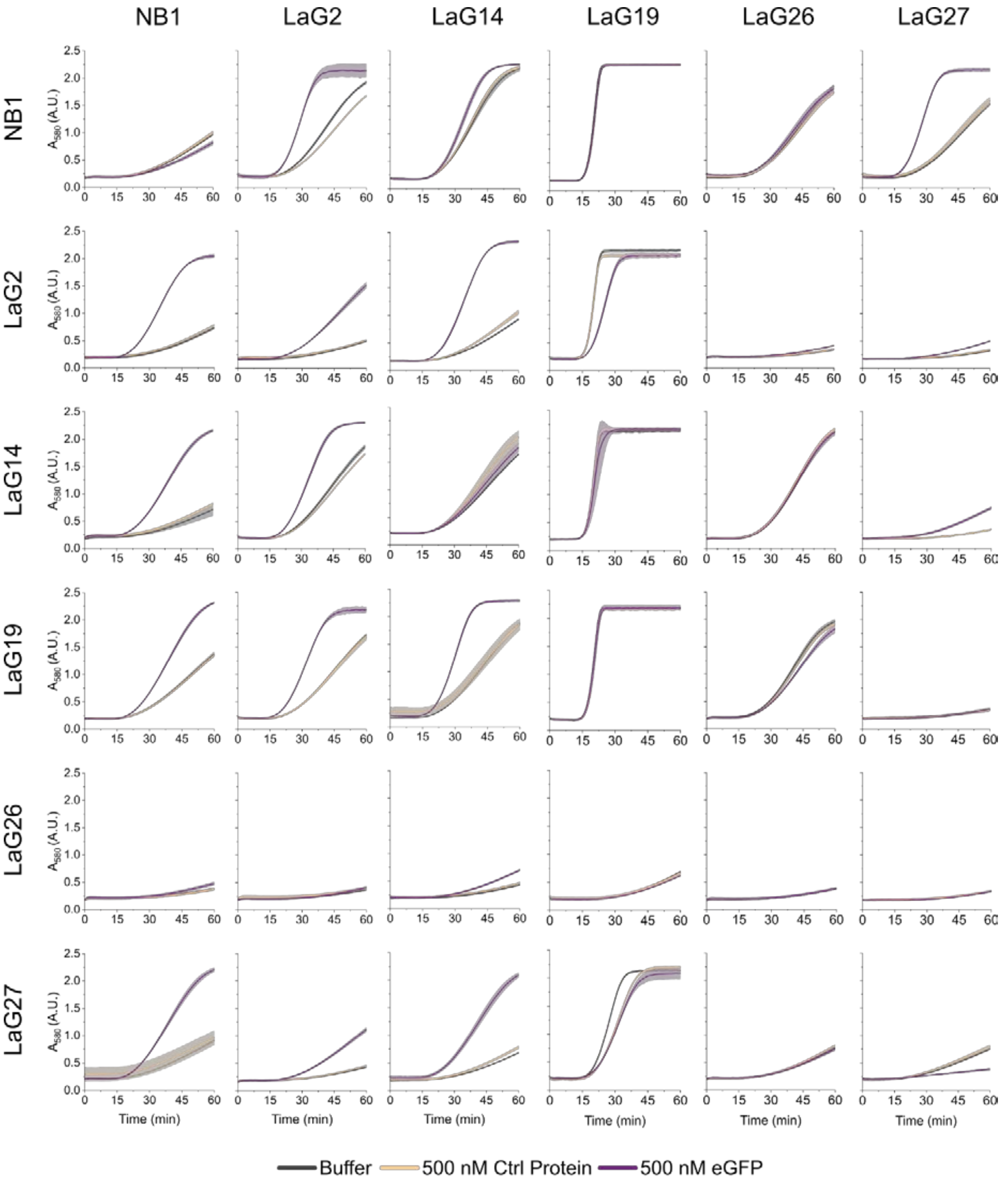


**Fig. S2.**

Absorbance data corresponding to the data in Figure 2A. Columns represent different T7RNAP$_{C}$ NB fusions and rows represent different T7RNAP$_{Nev}$ NB fusions. Lines and shaded areas represent the arithmetic mean ± standard deviation of n=3 technical replicates. Consistent with Figures 2B and 2C, the purple curve is for 500 nM eGFP, the yellow curve is for 500 nM control protein (mCherry), and the gray curve is for the buffer-only control.

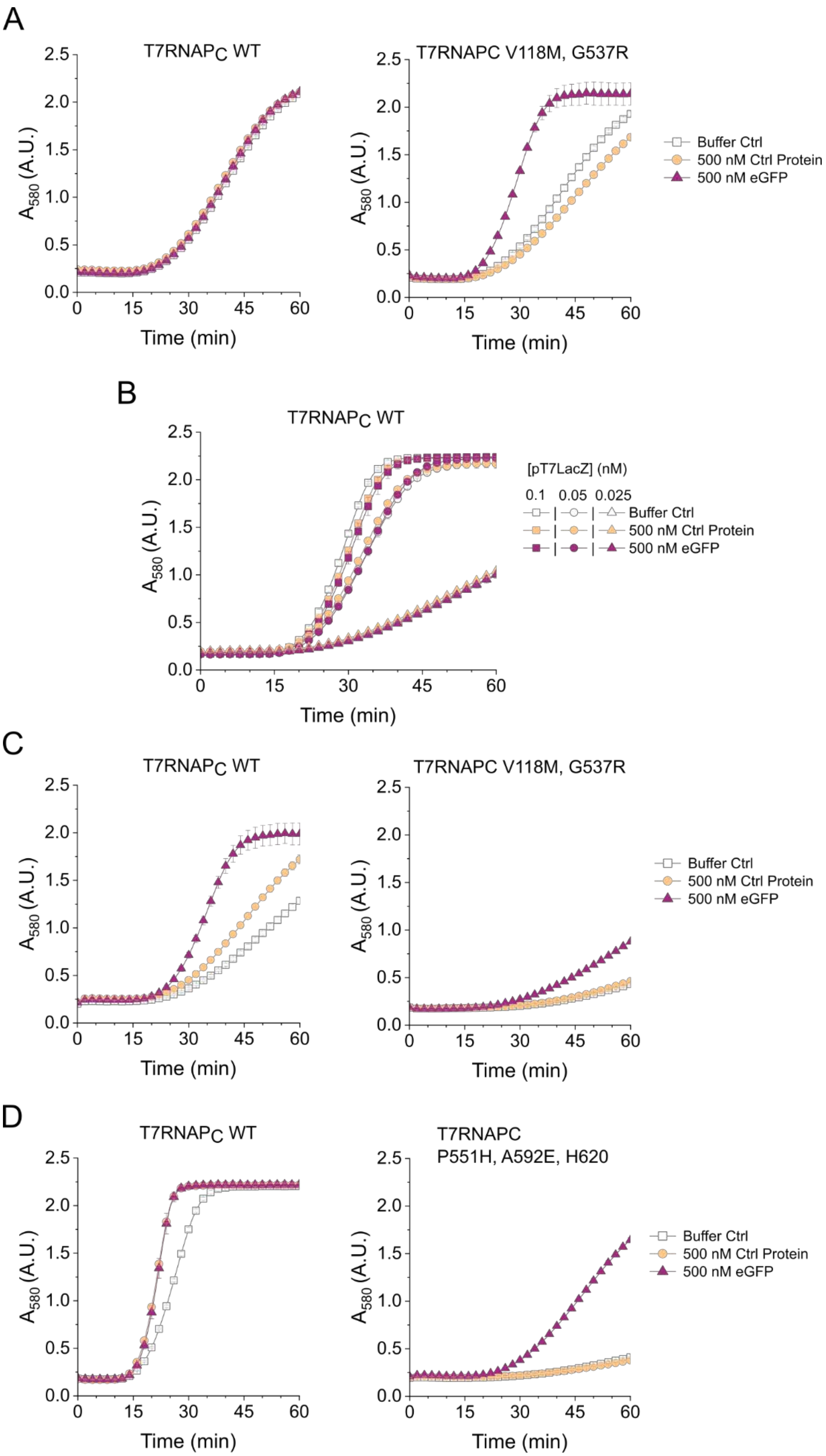
A
T7RNAPC WT
T7RNAPC V118M, G537R
A580 (A.U.)
Time (min)
Buffer Ctrl
500 nM Ctrl Protein
500 nM eGFP
B
T7RNAPC WT
[pT7LacZ] (nM)
0.1 0.05 0.025
Buffer Ctrl
500 nM Ctrl Protein
500 nM eGFP
C
T7RNAPC WT
T7RNAPC V118M, G537R
D
T7RNAPC WT
T7RNAPC
P551H, A592E, H620

**Fig. S3.**

TLISA performance when using either the WT T7RNAPC or the identified T7RNAPC mutations. (A) $T7RNAP_{Nev}$-NB1/LaG2-$T7RNAP_C$ eGFP TLISA with either (left) WT $T7RNAP_C$ or (right) mutated $T7RNAP_C$. (B) $T7RNAP_{Nev}$-NB1/LaG2-$T7RNAP_C$ eGFP TLISA using the WT $T7RNAP_C$ with a range of different pT7LacZ reporter plasmid concentrations. (C) $T7RNAP_{Nev}$-LaG2/GS2-$T7RNAP_C$ eGFP TLISA with either (left) WT $T7RNAP_C$ or (right) mutated $T7RNAP_C$. (D) $T7RNAP_{Nev}$-LaG2/3G86.32-$T7RNAP_C$ eGFP TLISA with either (left) WT $T7RNAP_C$ or (right) mutated $T7RNAP_C$.

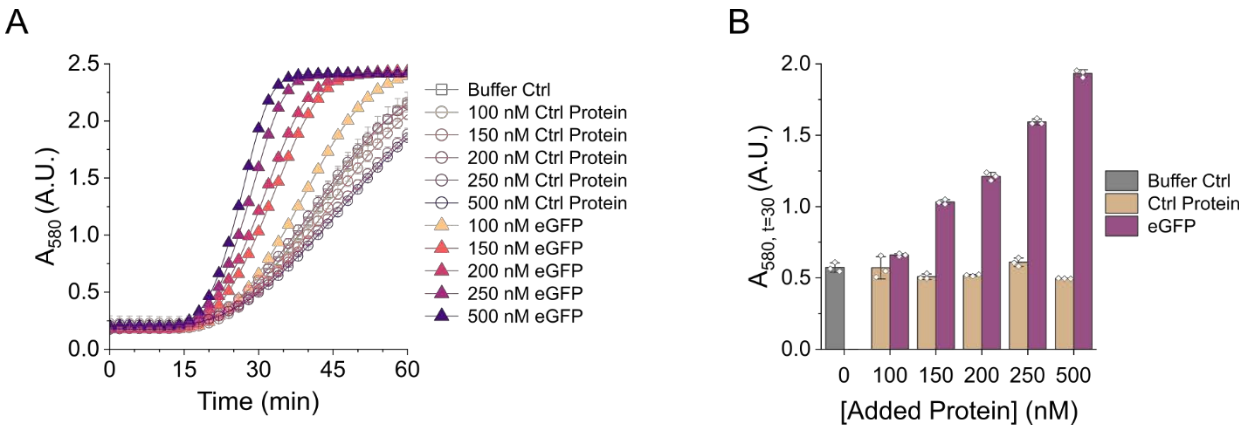


**Fig. S4.**

Detection range and LOD of the $T7RNAP_{Nev}$-LaG2/LaG14-$T7RNAP_{C}$ TLISA. (A) Absorbance data with increasing concentrations of eGFP or control protein showing an LOD of 100 nM eGFP after one hour. (B) Absorbance values at 30 minutes showing detection range. All reactions had 0.1 nM pT7LacZ. mCherry was used as the control protein. Bars represent the arithmetic mean ± standard deviation of n=3 technical replicates (white diamonds).

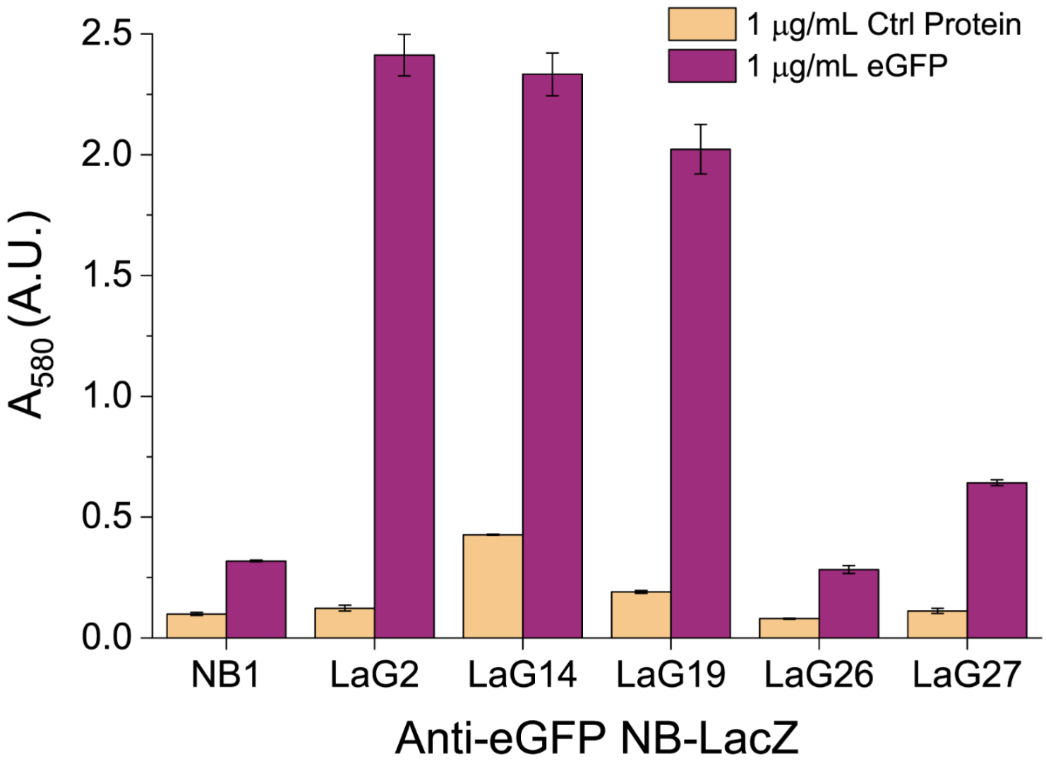


**Fig. S5.**

Direct ELISA validation of anti-eGFP NB expression and functionality in a CFE system. Anti-eGFP NBs were translationally fused to LacZ. TTR was used as the control protein. Bars and error bars represent the arithmetic mean ± standard deviation of n=3 technical replicates.

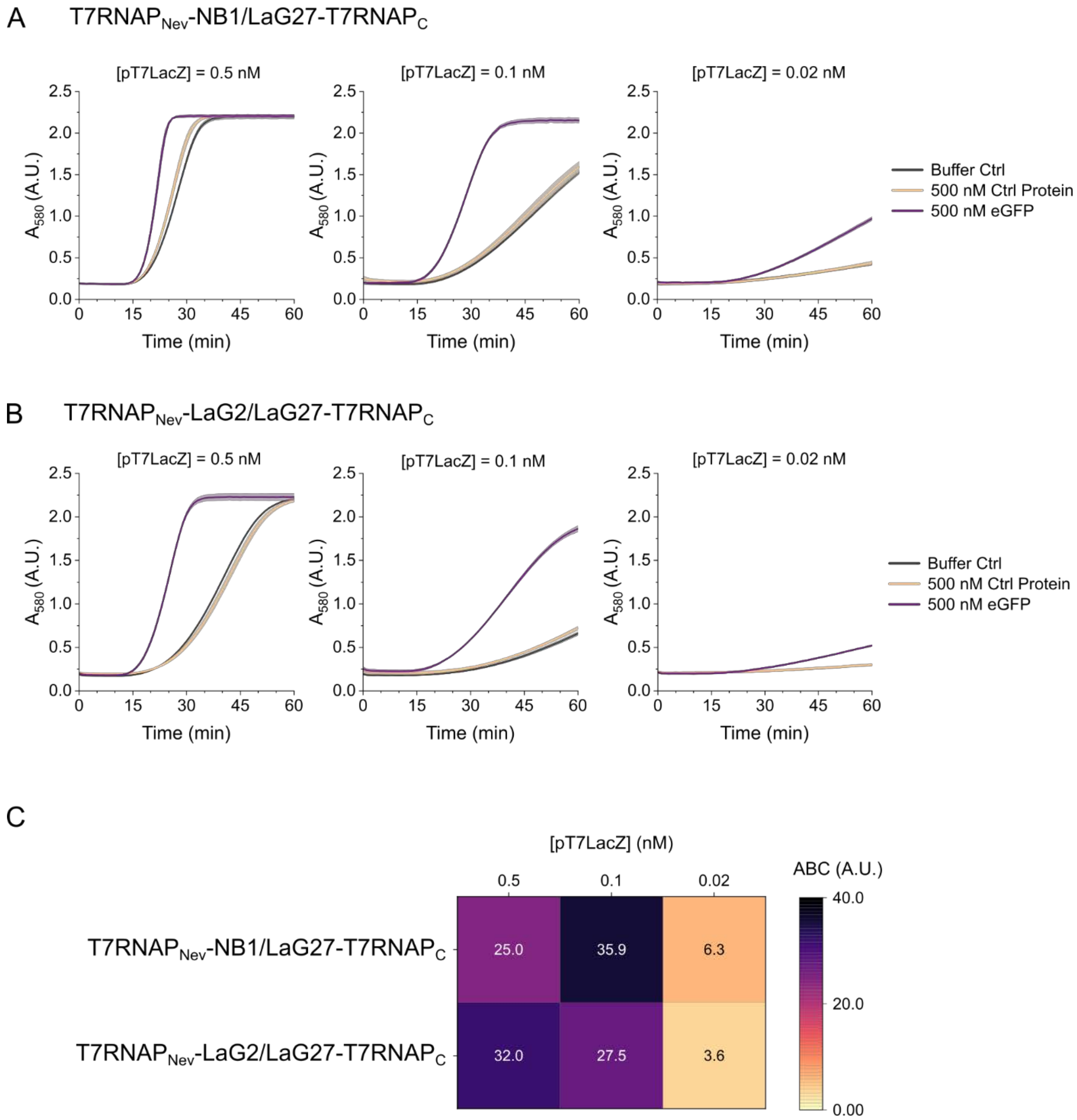


**Fig. S6.**

Optimal reporter plasmid concentration can vary across sensors. Absorbance data for (A) $T7RNAP_{Nev}$-NB1/LaG27-$T7RNAP_C$ and (B) $T7RNAP_{Nev}$-LaG2/LaG27-$T7RNAP_C$ with either 0.5 nM, 0.1 nM, or 0.02 nM pT7LacZ reporter plasmid. (C) Heatmap of ABC values for both sensors at all reporter plasmid concentrations. Shaded areas represent the standard deviation of the mean of n=3 technical replicates.

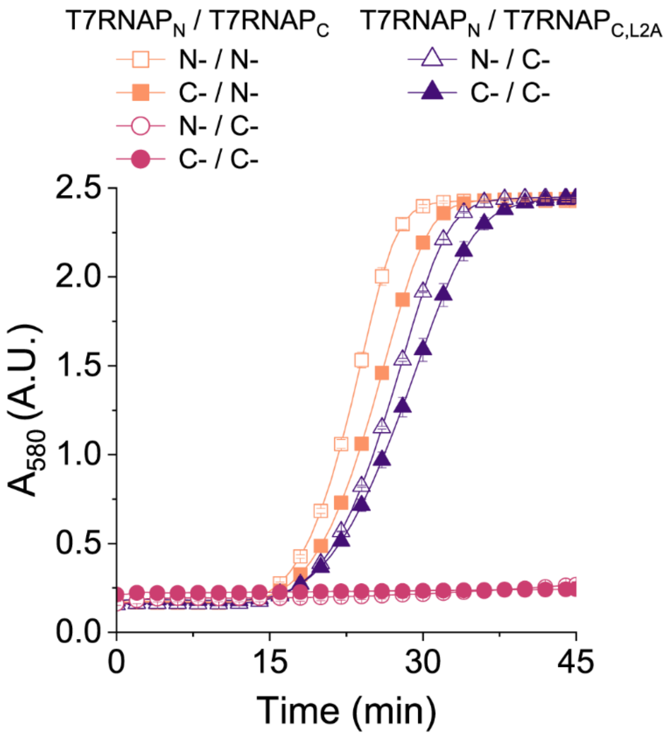


**Fig. S7.**

Spontaneous reassembly of T7RNAP fragments with NBs fused to different termini. All $T7RNAP_N$ fragments here are wild type to enable spontaneous reassembly. The evolved $T7RNAP_{C,L2A}$ variant enables C-terminal fusions. All reactions contained 0.1 nM pT7LacZ. Symbols represent the arithmetic mean ± standard deviation of n=3 technical replicates.

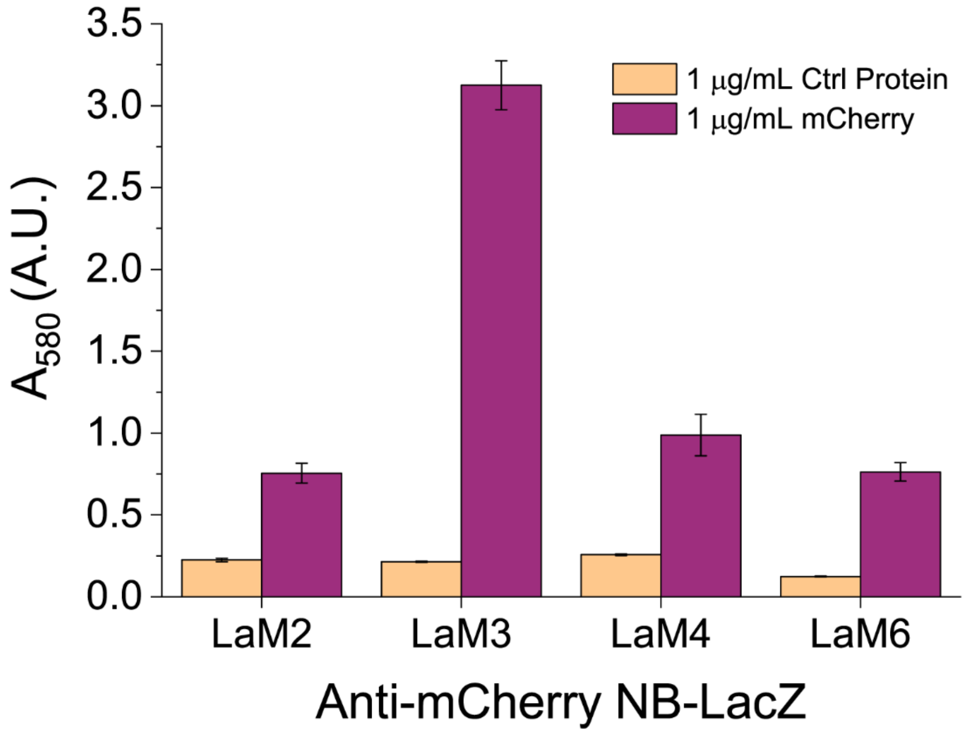


**Fig. S8.**

Direct ELISA validation of anti-mCherry NB expression in a CFE system. TTR was used as the control protein. Bars and error bars represent the arithmetic mean ± standard deviation of n=3 technical replicates.

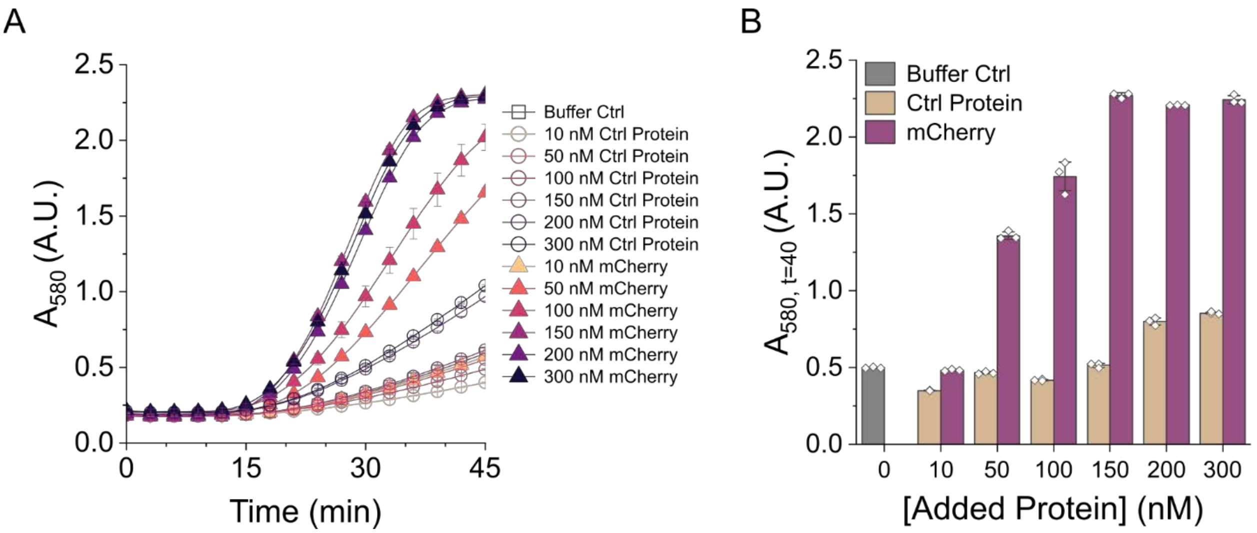


**Fig. S9.**

Detection range and LOD of the T7RNAP$_{Nev}$-LaM4/LaM2-T7RNAP$_{C}$ TLISA. (A) Absorbance data with increasing concentrations of mCherry or control protein showing an LOD of 50 nM mCherry. (B) Absorbance values at 40 minutes showing the detection range. All reactions had 0.05 nM pT7LacZ. TTR was used as the control protein. Bars represent the arithmetic mean ± standard deviation of n=3 technical replicates (white diamonds).

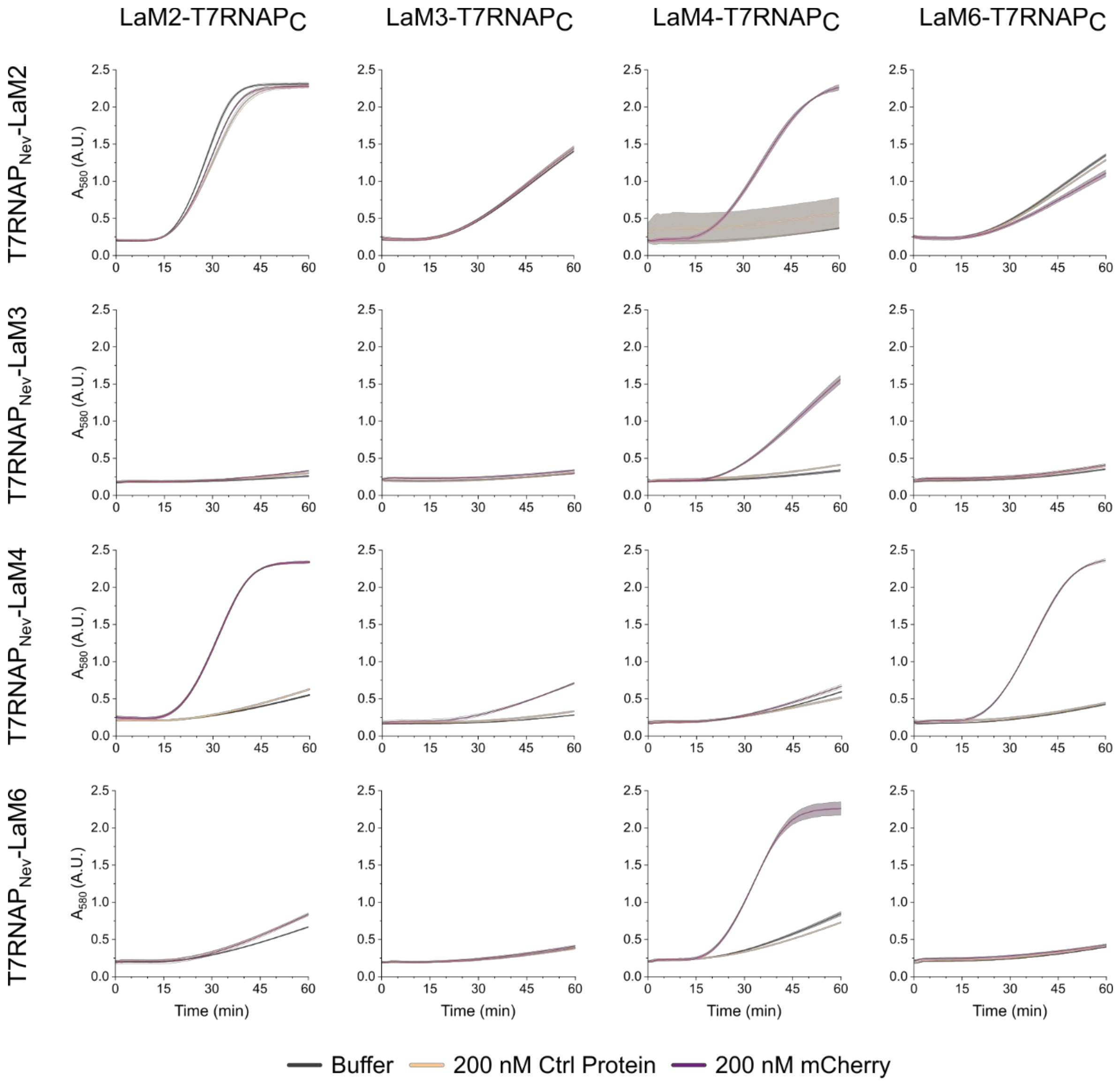


**Fig. S10.**

Absorbance data corresponding to the data in Figure 4B. Columns represent different T7RNAP$_C$ NB fusions and row represent different T7RNAP$_{Nev}$ NB fusions. Shaded areas represent the standard deviation of the mean of n=3 technical replicates. Consistent with Figure 4A, the purple curve is for 200 nM mCherry, the yellow curve is for 200 nM control protein (eGFP), and the gray curve is for the buffer-only control.

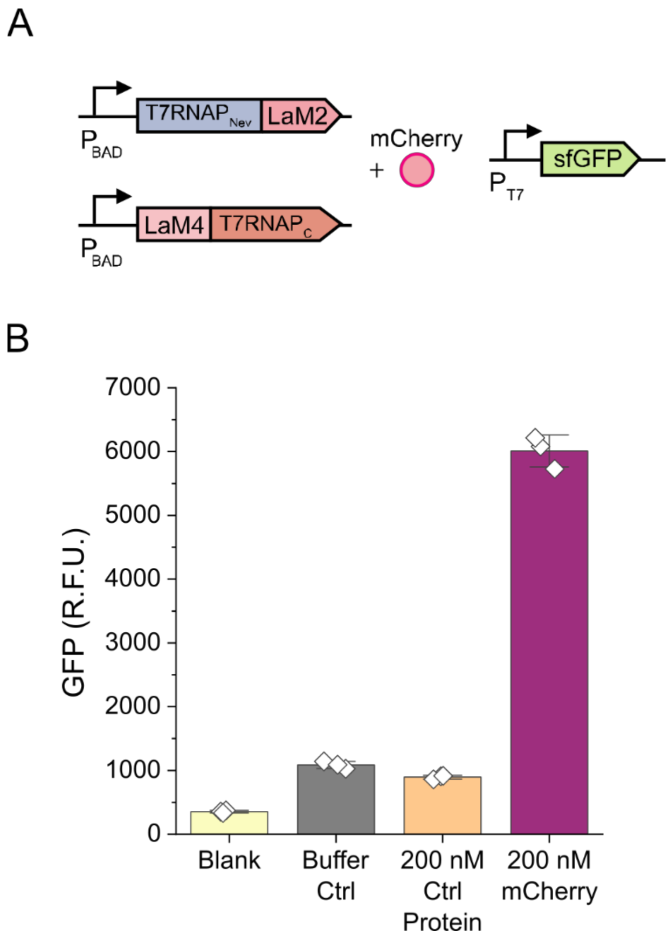


**Fig. S11.**

TLISA allows for different protein outputs. (A) Sensor circuit for $T7RNAP_{Nev}$-LaM2/LaM4-$T7RNAP_C$ mCherry TLISA using sfGFP as a protein reporter. (B) Fluorescent measurements (ex. 485 nm, em. 510 nm) after 3 hours of incubation at 37 °C. Blank represents the background fluorescence of the cell-free reaction and contains everything in the buffer control condition expect pT7sfGFP plasmid. 200 nM TTR was used as the control protein. Bars represent the arithmetic mean ± standard deviation of n=3 technical replicates (white diamonds).

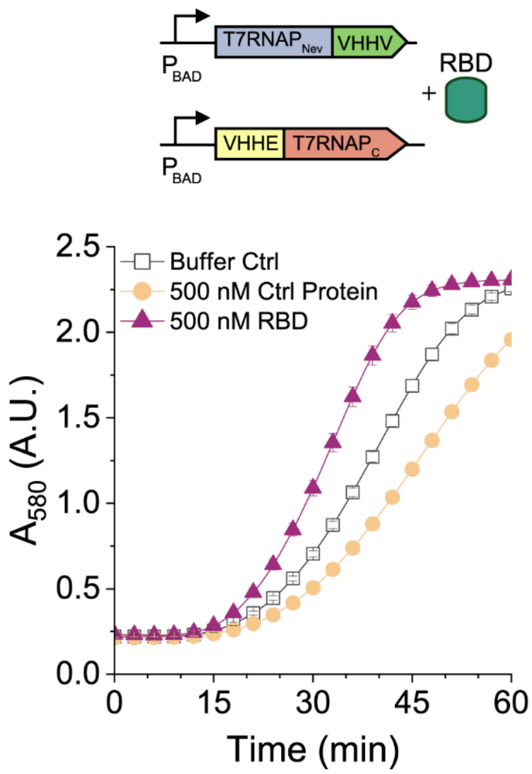


**Fig. S12.**

T7RNAPNev-VHHV/VHHE-T7RNAPC SARS-CoV-2 RBD TLISA. 500 nM mCherry was used as the control protein for all experiments. Symbols represent the arithmetic mean ± standard deviation of n=3 technical replicates.

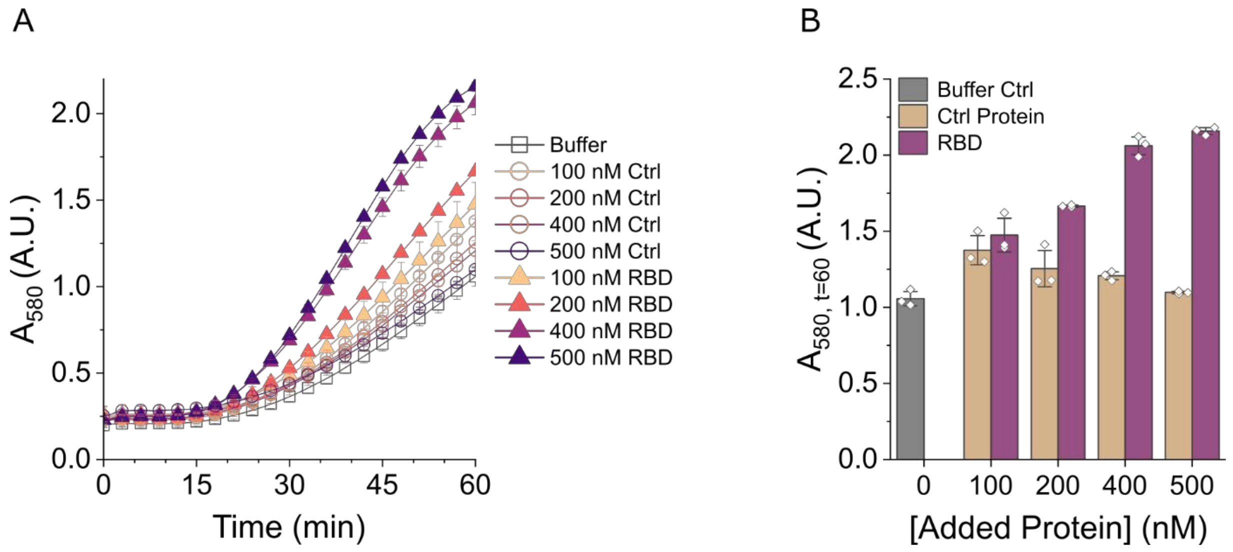


**Fig. S13.**

Determining the detection range and LOD of the T7RNAP$_{Nev}$-FSR22/VHHV-T7RNAP$_{C}$ biosensor for SARS-CoV-2 RBD. (A) Absorbance data with increasing concentrations of RBD or control protein showing an LOD of 200 nM RBD. (B) Absorbance values after 60 minutes of incubation at 37 °C. All reactions had 0.1 nM pT7LacZ. mCherry was used as the control protein. Bars represent the arithmetic mean ± standard deviation of n=3 technical replicates (white diamonds).

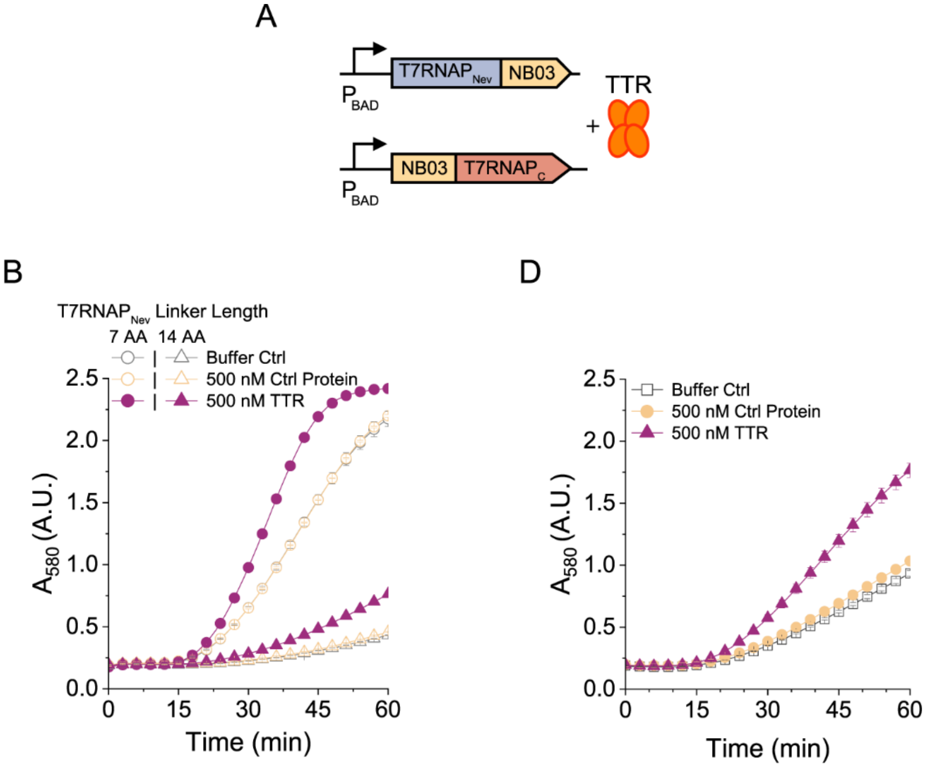


C

| Linker Length (AA) | ABC (A.U.) |
|---|---|
| 7 | 16.7577 |
| 14 | 6.7934 |

E

| [pT7LacZ] (nM) | ABC (A.U.) |
|---|---|
| 0.12 | 16.7577 |
| 0.05 | 15.8448 |

**Fig. S14.**

TTR sensor tuning. (A) Genetic sensing circuit for the TTR TLISA. (B) Absorbance data for TTR data with either a 7 AA linker (circles) or a 14 AA linker (triangles) on the T7RNAP$_{Nev}$-NB03 fragment with 0.12 nM pT7LacZ. (C) ABC values for the data in (B). (D) Absorbance data for the TTR sensor using the 7 AA linker on the T7RNAP$_{Nev}$-NB03 fragment with 0.05 nM pT7LacZ. (E) ABC values for 7 AA linker data in (B) and (D). 500 nM mCherry was used as the control protein for all experiments. Symbols represent the arithmetic mean ± standard deviation of n=3 technical replicates.

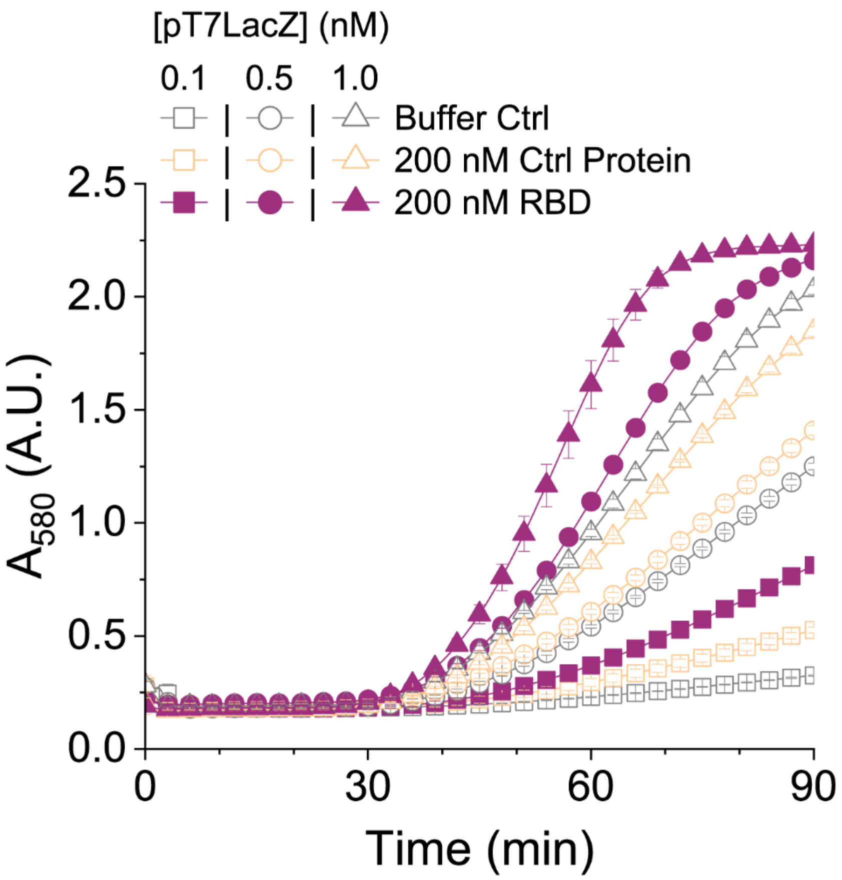


**Fig. S15.**

Increasing concentrations of pT7LacZ improves rate of reaction when sensing at room temperature. Absorbance data for T7RNAP$_{Nev}$-FSR22/VHHV-T7RNAP$_{C}$ SARS-CoV-2 RBD TLISA reaction in 20% v/v pooled human saliva incubated at 25 °C. 200 nM mCherry was used as the control protein. Symbols represent the arithmetic mean ± standard deviation of n=3 technical replicates.

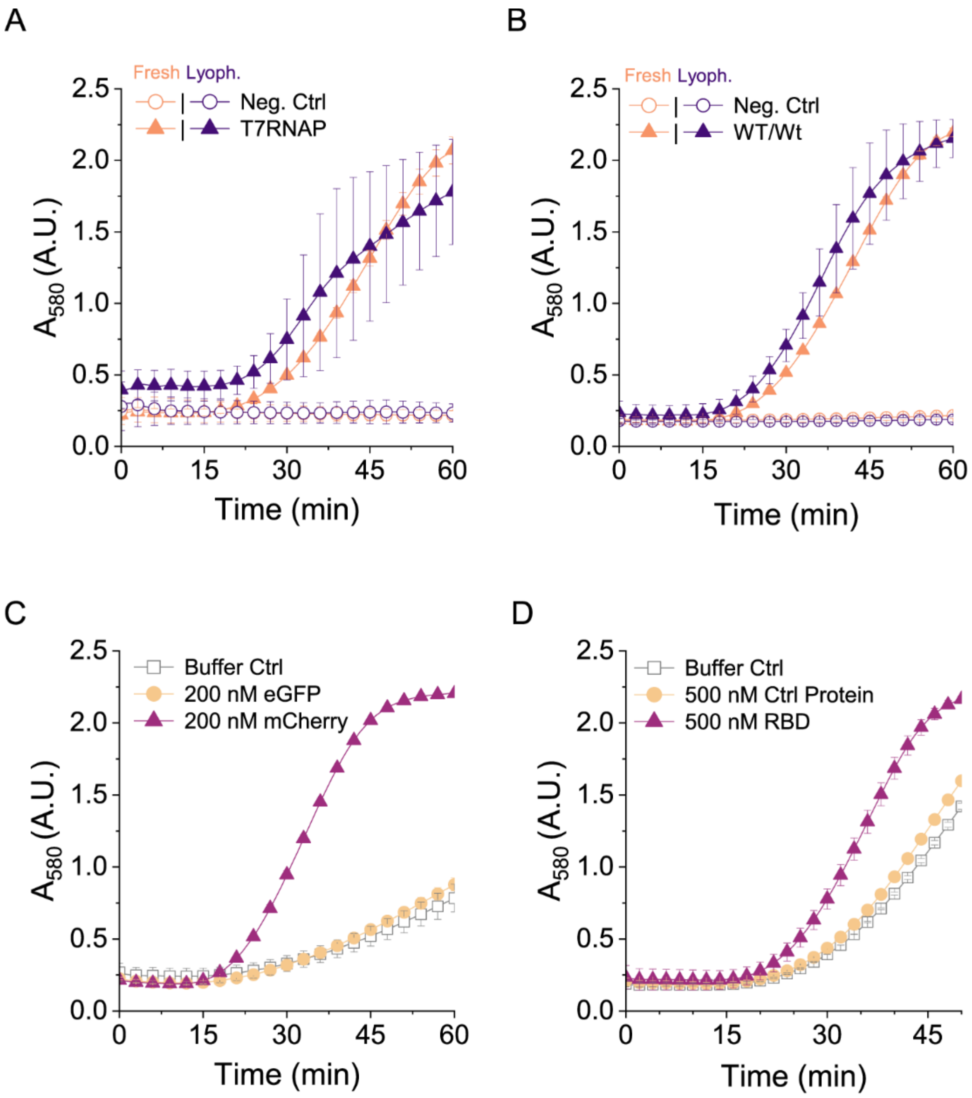


**Fig. S16.**

Successful lyophilization of cell-free reactions after 1 hour pre-expression reactions. (A) Lyophilized reactions after pre-expression of either a plasmid encoding full T7RNAP or an empty reaction (neg. ctrl) and rehydrated with 0.05 nM pT7LacZ. (B) Lyophilized reactions after pre-expression of plasmids encoding WT split T7RNAP fragments or an empty reaction (neg. ctrl) and rehydrated with 0.05 nM pT7LacZ. (C) Lyophilized T7RNAPNev-LaM4/LaM2-T7RNAPC mCherry TLISA rehydrated with 0.1 nM pT7LacZ. (D) Lyophilized T7RNAPNev-FSR22/VHHV-T7RNAPC SARS-CoV-2 RBD TLISA rehydrated with 0.5 nM pT7LacZ. Symbols represent the arithmetic mean ± standard deviation of n=3 technical replicates.

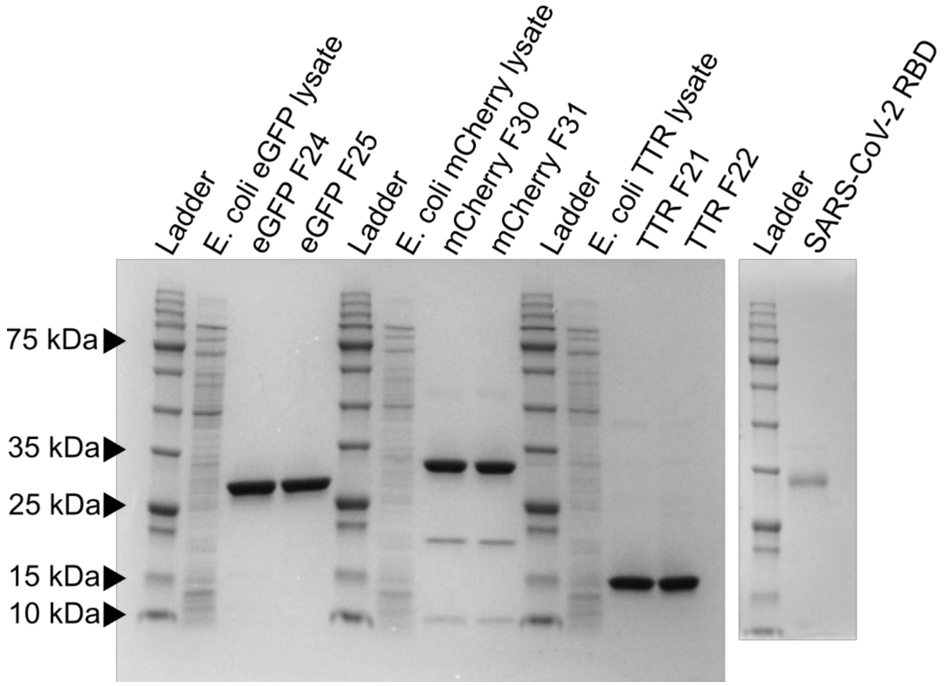


**Fig. S17.**

SDS-PAGE analysis of the purified proteins used in this study. The additional bands and high apparent MW of mCherry (Lanes F30 and F31) are expected and not evidence of impurities; the bands appearing at 20 kDa and 10 kDa are a result of mCherry fragmentation due to denaturing by boiling**(37,51)**.

| Plasmid | Mutation(s) | Figure(s) |
|---|---|---|
| NB1-T7RNAP$_C$ | V118M, G537R | 2A, S1 |
| LaG2-T7RNAP$_C$ | V118M, G537R | 1B, 2A, 3A-D, S1, S3, S5 |
| LaG14-T7RNAP$_C$ | V118M | 2A, S1, S6 |
| LaG27-T7RNAP$_C$ | V118M, G537R | 2A, S1, S4 |
| GS2-T7RNAP$_C$ | V118M, G537R | S3 |
| 3G86.32-T7RNAP$_C$ | P551H, A592E, H620N | 2C, S3 |
| NB1-Linker7-T7RNAP$_C$ | V118M, G537R | 3A |
| NB1-Linker28-T7RNAP$_C$ | V118M, G537R | 3A |
| LaM2-T7RNAP$_C$ | V118M | 4A, 4B, S9, S10 |
| LaM3-T7RNAP$_C$ | V118M | 4B, S10 |
| LaM4-T7RNAP$_C$ | V118M | 4B, S10, S11 |
| LaM6-T7RNAP$_C$ | V118M | 4B, S10 |
| VHHE-T7RNAP$_C$ | V118M, G537R | S12 |
| NB03-T7RNAP$_C$ | V118M | 5C, S14 |

**Table S1.**
Summary of plasmids containing mutations used in this study.

| Name | Type | Target antigen | Amino acid sequence | Figure(s) | Ref. |
|---|---|---|---|---|---|
| NB1 | Nanobody | eGFP | QVQLVESGGALVQPGGSLRLSCAASGFPVNRYSMRWYRQAPG KEREWVAGMSSAGDRSSYEDSVKGRFTISRDDARNTVYLQMN SLKPEDTAVYYCNVNVGFEYWGQGTQVTVSSK | 1B, 2A, 3A, 3B, 3C, 3D, S1-S3, S4A, S4C, S5 | (32) |
| LaG2 | Nanobody | eGFP | AQVQLVESGGGLVQAGGSLRLSCAASGRTFSNYAMGWFRQAP GKEREFVAAISWTGVSTYYADSVKGRFTISRDNDKNTVYVQM NSLIPEDTAIYYCAAVRARSFSDTYSRVNEYDYWGQGTQVTV | 1B, 2, 3, S1, S2, S3, S4B, S4C, S5 S10 | (33) |
| LaG14 | Nanobody | eGFP | AQVQLVESGGGLVQAGGSLRLSCAASGRTYSISAMGWFRQAP GKEREFVAGISRSGGTTYYADPVKGRFTISRDNAKNTVYLQM NSLKPEDTAVYYCAARARGWTTFPAREIEYDYWGQGTQVTV | 2A, S2, S5, S6 | (33) |
| LaG19 | Nanobody | eGFP | AQVQLVESGGGLVQAGGSLRLSCAASGPTGAMAWFRQAPGKE REFVGGISRSGTDTYYVDSVKGRFTIDRDNAKNTVYLQMNSL KPEDTAVYYCAARRSQILFTSRTDYEFWGQGTQVTV | 2A, 3E, S2, S5 | (33) |
| LaG26 | Nanobody | eGFP | AQVQLVESGGGLVQAGASMRLSCAASGITFSLYHWVWFRQAA GREHEFVAGIIRSGGETLSADSVKDRFIISRDDAKNTLYLQM NMLQPEDTATYYCAATHRADWYSSAFREYIFRGQGTQVTVS | 2A, S2, S5 | (33) |
| LaG27 | Nanobody | eGFP | ADVQLVESGGGLVQAGGSLRLSCTASGLTISTYNIGWFRQAP GKEREFVGIIIRNGDTTYYADSVKGRFTISRDNAKNTVYLQM NSVKPADAAVYSCGATVRAGAAAEQYNSYIFRGQGTQVTV | 2A, S2, S4, S5 | (33) |
| GS2 | Monobody | eGFP | VSSVPTKLEVVAATPTSLLISWDAPAVTVDHYYITYGETGHY WYYQAFAVPGSKSTATISGLSPGVDYTITVYAPFSVPVMSPI SINYRT | 2B, S3 | (34) |
| 3G86.32 | DARPin | eGFP | RGSGSDLGKKLLEAARAGQDDEVRILMANGADVNALDRFGLT PLHLAAQRGHLEIVEVLLKCGADVNAADLWGQTPLHLAATAG HLEIVEVLLKYGADVNALDLIGKTPLHLTAIDGHLEIVEVLL KHGADVNAQDKFGKTAFDISIDNGNEDLAEILQKLN | 2C, S3 | (52) |
| LaM2 | Nanobody | mCherry | AQVQLVESGGGLVQAGGSLRLSCATSGFTFSDYAMGWFRQAP GKEREFVAAISWSGHVTDYADSVKGRFTISRDNVKNTVYLQM NSLKPEDTAVYSCAAAKSGTWWYQRSENDFGSWGQGTQVTVS KEAI | 4, 6B, S9, S8, S10, S11, S16 | (33) |
| LaM3 | Nanobody | mCherry | AQVQLVQSGGGLVQAGGSLRLSCAASGRTFSDIAVGWFRQTP GKEREFVAAISWSGLIINYGDSVEDRFTISRDNAKSAVYLQM | 4B, S8, S10 | (33) |

| | | | | | |
|---|---|---|---|---|---|
| | | | NSLKPEDTAVYYCAARIGMNYYYAREIEYPYWGQGTQVTVSK CY | | |
| LaM4 | Nanobody | mCherry | AQVQLVESGGSLVQPGGSLRLSCAASGRFAESSSMGWFRQAP GKEREFVAAISWSGGATNYADSAKGRFTLSRDNTKNTVYLQM NSLKPDDTAVYYCAANLGNYISSNQRLYGYWGQGTQVTVSSP FT | 4, 6B, S9, S8, S10, S11, S16 | (33) |
| LaM6 | Nanobody | mCherry | AQVQLVESGGGLVQAGGSLRLSCVASGSAPSFFAMAWYRQSP GNERELVAALSSLGSTNYADSVKGRFTISMDNAKNTVYLQMN NVNAEDTAVYYCAAGDFHSCYARKSCDYWGQGTQVTVS | 4B, S8, S10 | (33) |
| VHHE | Nanobody | SARS CoV-2 RBD | QVQLVETGGGFVQPGGSLRLSCAASGVTLDYYAIGWFRQAPG KEREGVSCIGSSDGRTYYSDSVKGRFTISRDNAKNTVYLQMN SLKPEDTAVYYCALTVGTYYSGNYHYTCSDDMDYWGKGTQVT VSSGSGLNDIFEAQKIEWHE | 5A, S12 | (38) |
| VHHV | Nanobody | SARS CoV-2 RBD | QVQLVETGGGLVQPGGSLRLSCAASGFTFSSYAMGWARQVPG KGLEWVSYIYSDGSTEYQDSVKGRFTISRDNAKSTVYLQMNS LKPEDTAVYYCATEGSLGGWGRDFGSWGQGTQVTVSS | 5A, 5B, 6C, 6D, S12, S13, S15, S16 | (38) |
| FSR22 | DARPin | SARS CoV-2 RBD | GSSSSGMEQKLISEEDLDGYIPEAPRDGQAYVRKDGEWVLLS TFLGGGGSLQGGGGSLQGSDLGKKLLEAARAGQDDEVRILMA NGADVNACDPSGITPLHLAADKGHLEIVEVLLKYGADVNAMD VWGRTPLHLAAFTGHLEIVEVLLKYGADVNACDLNGYTPLHL AAGRGHLEIVEVLLKNGAGVNAQDKFGKTAFDISIDNGNEDL AEILQSSS | 5B, 6C, 6D, S13, S15, S16 | (39) |
| NB03 | Nanobody | TTR | QVQLQESGGGSVQAGGSLRLSCAASGNTYSYKVIGWFRQAPG KEREGIAAIYTGGVSSTRYADSVKGRFTISRDNTKNAVYLQM NSLKPEDTAMYYCAAGPLYDSTWFRAEKYNYWGQGTQVTVSS | 5C, S14 | (44) |

**Table S2.**

Sequences of nanobodies, monobodies, and DARPins used in this study.

| Name | Sequence | Figure(s) | Ref. |
|---|---|---|---|
| T7RNAP$_{N}$ | NTINIAKNDFSDIELAAIPFNTLADHYGERLAREQLALEHESYEMGEARFRKMFERQLKAG<br>EVADNAAAKPLITTLLPKMIARINDWFEEVKAKRGKRPTAFQFLQEIKPEAVAYITIKTTL<br>ACLTSADNTTVQAVASAIGRAIEDEARFGRIRDLEAKHFKKNVEEQLNKRVGHVYK | S7, S16 | (30) |
| T7RNAP$_{Nev}$ | NTINIAKNDFSDIELAAIPLNTLADHYGERSARGQLALEHESYEMGEARFRKMFECQLKAG<br>KVADNAAAKPLITTLLPKMIARINDWFEEVKAKRGRRPTAFKFLKEIKPEAVAYITIKTSL<br>ACLTSADNTTVQAVASAIGRTIEDEARFGRIRDLEAKHFKKNVEEQLNKRVGHVYK | 1-6, S1-S4, S6, S9-S16 | (31) |
| T7RNAP$_{C}$ | KAFMQVVEADMLSKGLLGGEAWSSWHKEDSIHVGVRCIEMLIESTGMVSLHRQNAGVVGQD<br>SETIELAPEYAEAIATRAGALAGISPMFQPCVVPPKPWTGITGGGYWANGRRPLALVRTHS<br>KKALMRYEDVYMPEVYKAINIAQNTAWKINKKVLAVANVITKWKHCPVEDIPAIEREELPM<br>KPEDIDMNPEALTAWKRAAAAVYRKDKARKSRRISLEFMLEQANKFANHKAIWFPYNMDWR<br>GRVYAVSMFNPQGNDMTKGLLTLAKGKPIGKEGYYWLKIHGANCAGVDKVPFPERIKFIEE<br>NHENIMACAKSPLENTWWAEQDSPFCFLAFCFEYAGVQHHGLSYNCSLPLAFDGSCSGIQH<br>FSAMLRDEVGGRAVNLLPSETVQDIYGIVAKKVNEILQADAINGTDNEVVTVTDENTGEIS<br>EKVKLGTKALAGQWLAYGVTRSVTKRSVMTLAYGSKEFGFRQQVLEDTIQPAIDSGKGLMF<br>TQPNQAAGYMAKLIWESVSVTVVAAVEAMNWLKSAAKLLAAEVKDKKTGEILRKRCAVHWV<br>TPDGFPVWQEYKKPIQTRLNLMFLGQFRLQPTINTNKDSEIDAHKQESGIAPNFVHSQDGS<br>HLRKTVVWAHEKYGIESFALIHDSFGTIPADAANLFKAVRETMVDTYESCDVLADFYDQFA<br>DQLHESQLDKMPALPAKGNLNLRDILESDFAFA | 1, 2, 3A, 3B, 3C, 4-6, S1-S4, S6, S7, S9-S16 | (30) |
| T7RNAP$_{C,L2A}$ | KAFMQAVEADMLSKGLLGGEAWSSWHKEDSIHVGVRCIEMLIESTGMVSLHRQNAGVVGQD<br>SETIELAPEYAEAIATRAGALAGISPMFQPCVVPPKPWTGITGGGYWANGRRPLALVRTHS<br>KKALMRYEDVYMPEVYKAINIAQNTAWKINKKVLAVANVITKWKNCPVEDIPAIEREELPM<br>KPEDIDTNPEALTAWKRAAAAVYRKDKARKSRRISLEFMLEQANKFANHKAIWFPYNMDWR<br>GRVYAVPMFNPQGNDMTKGLLTLAKGKPIGKEGYYWLKIHGANCAGVDKVPFPERIKFIEE<br>NHENIMACAKSPLGNTWWAEQDSPFCFLAFCFEYAGVQHHGLSYNCSLPLAYDESCSGIQH<br>FSAMLRDEVGGRAVNLIPSETVQDIYGIVAKKVNEILQADAINGTDNEVVTVTDENTGEIS<br>EKVKLGTKALAGQWLAYGVTRSVTKRSVMTLAYGSKEFGFRQQVLEDTIQPAIDSGKGLMF<br>TQPNHAAGYMAKLIWESASVTVVAAVEAMNWLKSAAKLLAAEVKDKKTGEILRKRCSVHWV<br>TPDGFPVWQEYKKPIQTRLNLMFLGQFRLQPTINTNKDSEIDAHKQESGIAPNFVHSQDGS<br>HLRKTVVWAHEKYGIESFALIHDSFGTIPADAANLFKAVRETMVDTYESCDVLADFYDQFA<br>DQLHESQLDKMPALPAKGNLNLRDILESDFAFA | 3D, S7 | (36) |

**Table S3.**
Sequences of the T7RNAP fragments used in this study.

| Name | Sequence | Figure(s) |
|---|---|---|
| Linker14 | GGSGSSGGSGSGSS | 1-6, S1-S4, S6, S7, S9-S15, S16C, S16D |
| Linker7 | GGSGSSG | 3A, 3B, S14 |
| Linker28 | GGSGSSGGSGSGSSGGSGSSGGSGSGSS | 3A, 3B |

**Table S4.**

Sequences of the flexible amino acid linkers used in this study.

| Name | Sequence |
|---|---|
| eGFP | 6xHis Tag-eGFP<br><br>MHHHHHHASKGEELFTGVVPILVELDGDVNGHKFSVSGEGEGDATYGKLTLKFICTTGKLPVPWPTLVTTLC<br>YGVQCFSRYPDHMKRHDFFKSAMPEGYVQERTIFFKDDGNYKTRAEVKFEGDTLVNRIELKGIDFKEDGNIL<br>GHKLEYNYNSHNVYIMADKQKNGIKVNFKTRHNIEDGSVQLADHYQQNTPIGDGPVLLPDNHYLSTQSALSK<br>DPNEKRDHMVLLEFVTAAGITHGMDELYN |
| mCherry | 6xHis Tag-mCherry<br><br>MHHHHHHVSKGEEDNMAIIKEFMRFKVHMEGSVNGHEFEIEGEGEGRPYEGTQTAKLKVTKGGPLPFAWDIL<br>SPQFMYGSKAYVKHPADIPDYLKLSFPEGFKWERVMNFEDGGVVTVTQDSSLQDGEFIYKVKLRGTNFPSDG<br>PVMQKKTMGWEASSERMYPEDGALKGEIKQRLKLKDGGHYDAEVKTTYKAKKPVQLPGAYNVNIKLDITSHN<br>EDYTIVEQYERAEGRHSTGGMDELYK |
| SARS-CoV2 RBD | Leader Peptide-RBD-Linker-SpyTag-6xHis Tag<br><br>METDTLLLWVLLLWVPGSTGDRVQPTESIVRFPNITNLCPFGEVFNATRFASVYAWNRKRISNCVADYSVLY<br>NSASFSTFKCYGVSPTKLNDLCFTNVYADSFVIRGDEVRQIAPGQTGKIADYNYKLPDDFTGCVIAWNSNNL<br>DSKVGGNYNYLYRLFRKSNLKPFERDISTEIYQAGSTPCNGVEGFNCYFPLQSYGFQPTNGVGYQPYRVVVL<br>SFELLHAPATVCGPKKSTNLVKNKCVNFGGSGGSAHIVMVDAYKPTKHHHHHH |
| Transthyretin (TTR) | 6xHis Tag-TTR Monomer<br><br>MHHHHHHGPTGTGESKCPLMVKVLDAVRGSPAINVAVHVFRKAADDTWEPFASGKTSESGELHGLTTEEQFV<br>EGIYKVEIDTKSYWKALGISPFHEHAEVVFTANDSGPRRYTIAALLSPYSYSTTAVVTNPKE |

**Table S5.**

Sequences of the protein antigens used in this study.
